# GATA4 regionalizes intestinal metabolism and barrier function to prevent immunopathology

**DOI:** 10.1101/2021.10.28.466194

**Authors:** Zachary M. Earley, Wioletta Lisicka, Joseph J. Sifakis, Raúl Aguirre-Gamboa, Anita Kowalczyk, Jacob T. Barlow, Dustin G. Shaw, Valentina Discepolo, Ineke L. Tan, Saideep Gona, Jordan D. Ernest, Polly Matzinger, Luis B. Barreiro, Andriy Morgun, Albert Bendelac, Rustem F. Ismagilov, Natalia Shulzhenko, Samantha J. Riesenfeld, Bana Jabri

**Affiliations:** Committee on Immunology, University of Chicago, Chicago, IL, USA; Department of Medicine, University of Chicago, Chicago, IL, USA; Department of Chemistry, University of Chicago, Chicago, IL, USA; Division of Biology and Biological Engineering, California Institute of Technology, CA, USA; Department of Medical Translational Sciences and European Laboratory for the Investigation of Food Induced Diseases, University of Federico II, Naples, IT; Department of Gastroenterology and Hepatology, University of Groningen and University of Medical Center Groningen, Groningen, NL; Genetics, Genomics, and Systems Biology, University of Chicago, Chicago, IL, USA; Ghost Lab. National Institute of Allergy and Infectious Diseases, National Institutes of Health, Bethesda, MD, USA; College of Pharmacy, Oregon State University, Corvallis, OR, USA; Department of Pathology, University of Chicago, Chicago, IL, USA; Division of Chemistry and Chemical Engineering, California Institute of Technology, CA, USA; Department of Biomedical Sciences, Oregon State University, Corvallis, OR, USA; Pritzker School of Molecular Engineering, University of Chicago, Chicago, IL, USA

## Abstract

Different regions of the gastrointestinal tract have distinct digestive and absorptive functions, which may be locally disrupted by infection or autoimmune disease. Yet, the mechanisms underlying intestinal regionalization and its dysregulation in disease are not well understood. Here, we used mouse models, transcriptomics, and immune profiling to show that regional epithelial expression of the transcription factor GATA4 prevented adherent bacterial colonization and inflammation in the proximal small intestine by regulating retinol metabolism and luminal IgA. Loss of epithelial GATA4 expression increased mortality in mice infected with *Citrobacter rodentium*. In active celiac patients with villous atrophy, low GATA4 expression was associated with metabolic alterations, mucosal *Actinobacillus*, and increased IL-17 immunity. This study reveals broad impacts of GATA4-regulated intestinal regionalization and highlights an elaborate interdependence of intestinal metabolism, immunity, and microbiota in homeostasis and disease.

**One-sentence summary:** Epithelial GATA4 regulates intestinal regionalization of bacterial colonization, metabolic pathways, and tissue immunity.

## Introduction

Each region of the gastrointestinal tract plays a distinct physiological role, featuring specific metabolic functions, host-microbe interactions, and immune responses. The duodenum and jejunum of the proximal intestine are optimized to digest and absorb critical nutrients. Despite having a large surface area in close contact with luminal contents, these regions also exhibit relatively low microbial growth and adhesion (*1*). In contrast, the ileum of the distal small intestine has shorter villi, less optimized for nutrient uptake, and instead specializes in reabsorption of bile acids and vitamin B12 (*1*). The ileum harbors more adherent commensal microbes, such as *segmented filamentous bacteria* (SFB), which promote local T_H_17 differentiation and inflammatory IL-17 responses and protect the host from enteropathogenic infections (*2–4*). Finally, the colon is characterized by a thick mucus layer that permits absorption of water, electrolytes, and numerous microbial metabolites, while preventing the microbiota from invading the intestinal epithelium and driving inflammatory immunity (*1, 5, 6*). Together, these patterns suggest that each region uniquely balances its critical digestive and barrier immune functions, yet the regulatory mechanisms controlling regionalization remain elusive.

One known regulator of intestinal regionalization is the transcription factor GATA4, which is selectively expressed in the intestinal epithelial cells (IECs) of the duodenum and jejunum (*7, 8*). Deleting GATA4 in IECs was shown to alter lipid absorption in the jejunum and drive major transcriptional changes, including down-regulation of jejunal-specific genes and up-regulation of ileal-specific genes (*8*). These results suggest that epithelial GATA4 has a profound role controlling numerous cell types in the tissue, a regulatory circuit that has not yet been dissected.

Using GATA4-deficient mice, we investigated how GATA4 controls barrier function and bacterial colonization in the jejunum through its key metabolic functions. We also assessed the impact of disrupted intestinal regionalization on host susceptibility to pathology, both in the context of enteric bacterial infection and human celiac disease, a dietary-induced autoimmune-like disorder of the proximal small intestine.

## Results

### GATA4 regulates regionalization of intestinal immunity in a microbiota-dependent manner

Analysis of cytokine production in T cells from the intestinal track of specific-pathogen-free (SPF) and germ-free (GF) mice, revealed regional-specific immune regulation (Fig. 1A). For instance, despite the microbial load being considerably higher in the colon than in the ileum, inflammatory microbiota-dependent IFN-γ^+^ and IL-17^+^T cell responses were significantly higher in the ileum. In contrast, regulatory IL-10^+^ T cell responses were found at similar levels in the ileum and colon, and were dependent on the presence of the microbiota in both locations. Furthermore, the microbiota was required for the presence of intraepithelial IFN-γ^+^ CD4+ T cells in the ileum but not the proximal small intestine.

**Figure 1.**
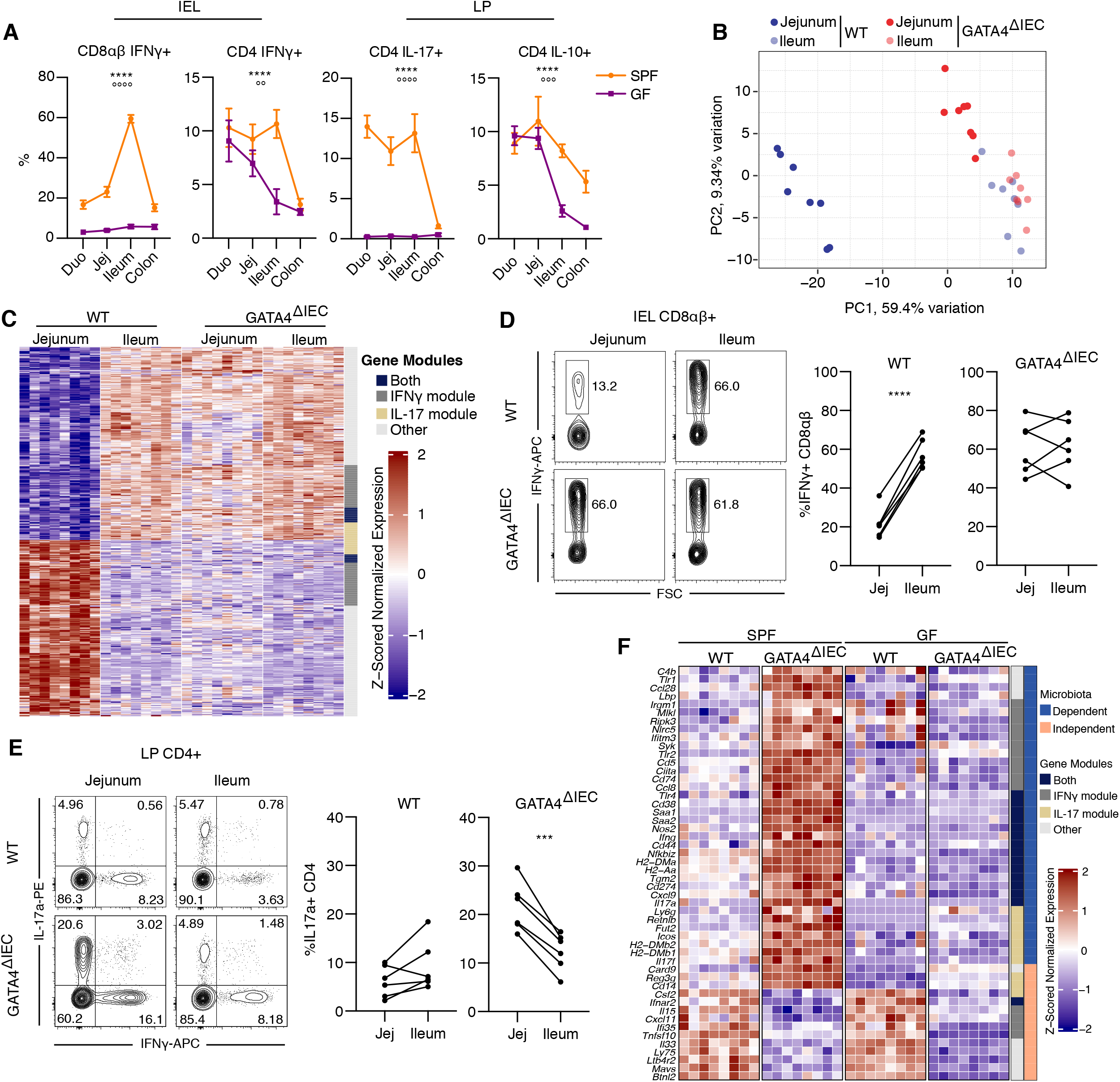
Intestinal epithelial cells control microbiota-dependent and independent regionalization of tissue immunity through GATA4. **(A)** Percentage (y axis) of cells stained for cytokines IFNγ, IL-17, or IL-10 (panel titles), as measured by intracellular flow cytometry, among all CD8αβ (leftmost panel) or CD4 (other panels) T cells isolated from intraepithelial lymphocytes (IELs, left set) or the lamina propria (LP, right set) of each intestinal segment (x axis) of SPF (orange circle) or GF (purple square) mice and then stimulated by phorbol myristate acetate (PMA) and ionomycin. ^****^, P<0.0001, effect due to region; ^oooo^ P<0.0001, ^ooo^ P<0.001, ^oo^ P< 0.01, effect due to microbiota; two-way ANOVA of microbiota and region impact on cytokine levels *N*= 5-7 mice/group. **(B)** Top two principal components (PCs) (x, y axes) of the expression of region-specific, GATA4-regulated genes, based on RNA-seq of tissue samples (dots) from the jejunum (dark colors) and ileum (light colors) of WT (blues) and GATA4^ΔIEC^ (reds) mice. *N=* 8 mice/group. **(C)** Heatmap of the z-scored expression (color) in tissue samples of region-specific, GATA4-regulated immune genes (rows) from the jejunum and ileum (columns) of WT (left set) and GATA4^ΔIEC^ (right set) mice. Of 625 total genes, 145 are in only the IFNγ module, 54 in only the IL-17 module, 39 in both modules, and 387 in neither (annotation column) (Methods). *N*= 8 mice/group. **(D)** Representative (left) and summary (right) plots of the frequencies of IFN-γ^+^ cells among CD8αβ T cells in the IELs of the jejunum and ileum of WT and GATA4^ΔIEC^ mice. **** P<0.0001, paired t-test. *N*= 6 mice/group. **(E)** Representative (left) and summary (right) plots of the frequencies of IL-17a^+^ cells among CD4 T cells in the LP of the jejunum and ileum of WT and GATA4^ΔIEC^ mice. *** P<0.001, paired t-test. *N*= 6 mice/group **(F)** Heatmap of the z-scored expression (color) in tissue samples of 50 selected microbiota dependent and independent (right annotation column), region-specific, GATA4-regulated immune genes (rows) in the jejunum (columns) of SPF (left set) and GF (right set) WT and GATA4^ΔIEC^ mice. Gene modules (left annotation column) as in C.

To determine how IECs shape regional tissue immunity, we analyzed mice in which GATA4 was selectively depleted in IECs (GATA4fl/fl villin-cre+ or GATA4^ΔIEC^) (*8*). First, we confirmed that GATA4 expression was restricted to duodenal and jejunal IECs, and that, in its absence, jejunal IECs acquired an ileal-like transcriptional program (S1A, B). GATA4 strongly repressed ileal-specific genes (*Fabp6, Slc10a2*) involved in the enterohepatic circulation of bile acids (Fig. S1C). In contrast, GATA4 induced expression of lipid metabolic genes in the jejunum involved in retinol metabolism (*Adh1*), and fat digestion and absorption (*Cd36, Fabp1, Dgat2, Apoa4)*. GATA4 also played an important role in controlling jejunal epithelial uptake of vitamins and folate (Slc46a1, Pdxk) (Fig. S1C). To systematically assess the impact of GATA4 on regional tissue functions, we performed total tissue RNA sequencing in GATA4^ΔIEC^ and littermate control wild-type (WT) mice (GATA4fl/fl villin-cre-). Our analysis identified what we term “region-specific, GATA4-regulated” genes, which are defined by expression that distinguishes “ileum-like” tissue, *i.e*., WT ileum and GATA4^ΔIEC^ jejunum, from WT jejunum (Fig. 1B, C). Intersecting region-specific, GATA4-regulated genes with public databases of immune genes (*9*) revealed distinct immune signatures of WT jejunum and ileum, a regionalization of tissue immunity that was lost in GATA4^ΔIEC^ mice (Fig. 1B). Indeed, among the 625 regional-specific, GATA4-regulated immune genes, more than a third (238) were published potential targets of IFNγ or IL-17 regulation (Fig. 1C).

Consistent with GATA4-regulated regionalization of IFNγ and IL-17 immune pathways, in the absence of epithelial GATA4, the frequency of intraepithelial IFNγ^+^ cells among CD8αβ T cells in the jejunum increased to the levels observed in the ileum (Fig. 1D). Furthermore, GATA4 deficiency led to a heightened T_H_17 response in the jejunum, with frequencies of IL-17^+^ cells among CD4 T cells surpassing those in the ileum (Fig. 1E). The heightened IFNγ and IL-17 immune responses observed in the jejunum of GATA4^ΔIEC^ mice were generally microbiota dependent (Fig. 1F, S1D-F). Intriguingly, though, some particular immune genes, such as *Csf2*, which encodes for GM-CSF, the anti-viral response genes *Ifnar2* and *Mavs*, and the tissue alarmins IL-33 and IL-15, were GATA4 regulated in a microbiota-independent manner (Figure 1F). Among epithelial, region-specific GATA4-regulated genes, the microbiota-independent subset was enriched, compared to the microbiota-dependent subset, in direct targets of GATA4 (Fig. S1G), as indicated by GATA4 binding of promoter regions in published ChIP-seq data (hypergeometric distribution test; p-value = 3.21E-80) (Fig. S1G) (*10, 11*). Taken together, these results demonstrate that GATA4 in IECs regulated regional tissue immunity and blocked development of microbiota-dependent inflammatory immune responses in the proximal intestine.

### GATA4 shapes regional immunity by controlling colonization of *segmented filamentous bacteria*

To investigate which specific microbiota may trigger inflammatory immune responses in the absence of GATA4 in the proximal small intestine, we performed 16S ribosomal RNA sequencing of luminal- and mucosal-associated bacterial communities in the jejunum and ileum of littermate control WT and GATA4^ΔIEC^ mice. This analysis revealed that in the jejunum, SFB (*Candidatus arthromitus*) loads were much higher in GATA4^ΔIEC^ relative to WT mice (Fig. 2A, S2A). Furthermore, GATA4 was required to prevent SFB from colonizing (Fig. 2A, S2A) and adhering (Fig. 2B) to IECs in the jejunum. In contrast, and in accordance with the lack of GATA4 expression in the ileum, no changes in bacterial composition were observed in the ileum of GATA4^ΔIEC^ versus WT mice (Fig. S1A).

**Figure 2.**
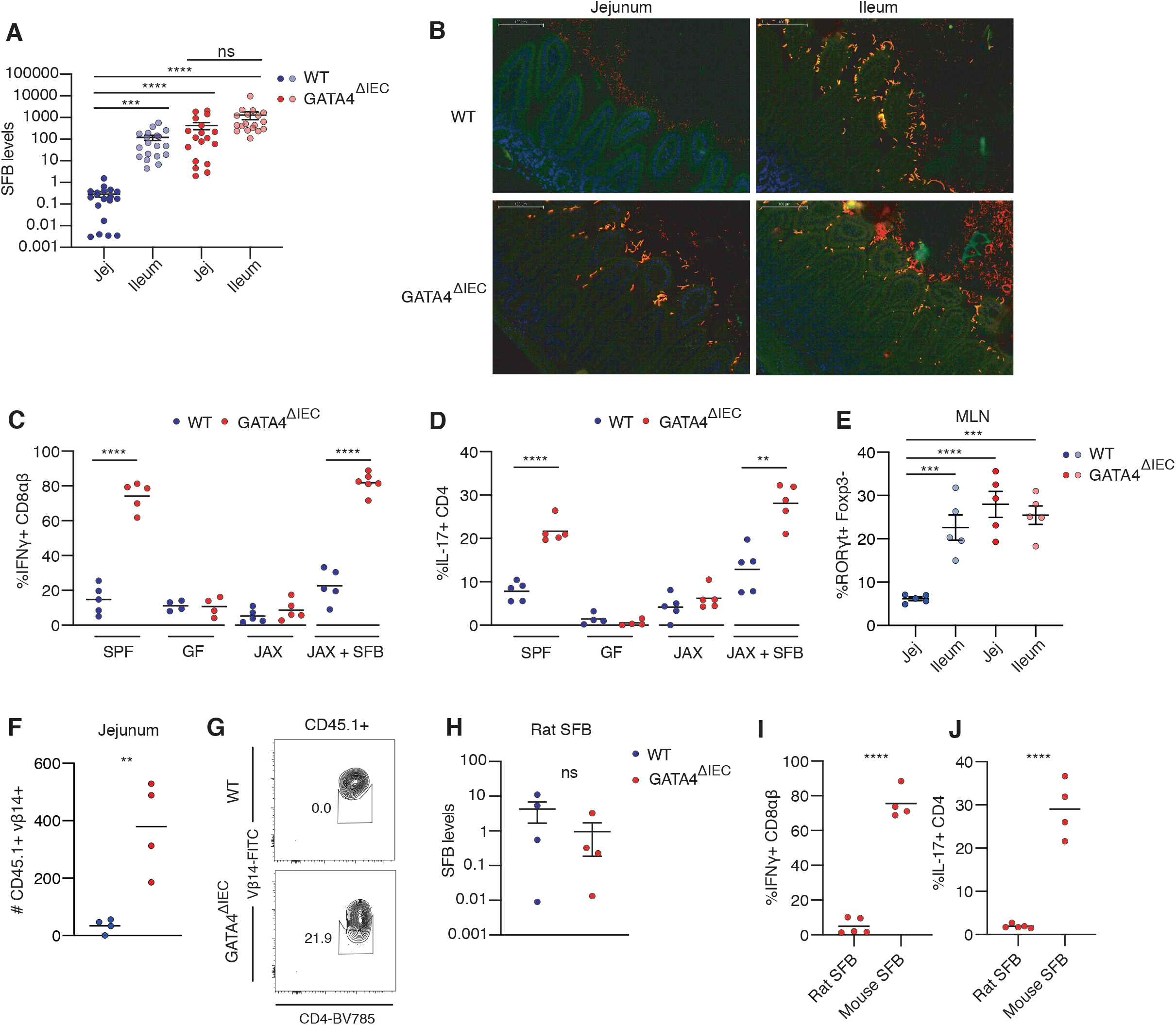
GATA4 controls the colonization of *segmented filamentous bacteria* to shape tissue immunity. **(A)** SFB load (y axis), as measured by qPCR, relative to the amount of host DNA in mucosal scrapings (dots) of jejunum (dark colors) and ileum (light colors) from WT (blues) and GATA4^ΔIEC^ (reds) mice. **** P<0.0001, *** Pp<0.001, ** P<0.01,Kruskal-Wallis with Dunn multiple comparison test. *N*= 18-19 mice/group. **(B)** Representative FISH staining of jejunal (left) and ileal (right) tissue of WT (top) and GATA4^ΔIEC^ (bottom) mice using universal 16s rRNA probes (Alexa 546) and a probe specific for SFB 16s (Alexa 488) and counterstained with DAPI. The overlay of the 16s probes represents SFB. (**C** and **D**) Frequency (y axis) of IFNγ^+^ cells among CD8αβ T cells (C) or IL-17a^+^ cells among CD4 T cells (D) in jejunum samples (dots) from SPF, GF, Jackson (JAX) microbiota transfer, and Jackson microbiota + SFB transfer (x axis) into WT and GATA4^ΔIEC^ mice (color, as in A). **** P<0.0001, ** P<0.01, t-test. *N*= 4-6 mice/group. **(E)** Frequency (y axis) of RORγt^+^ FOXP3^-^ cells among transferred CD45.1^+^ CD4^+^ Vβ14^+^ 7B8^+^ T cells in the MLN (dots) draining the jejunum and ileum of CD45.2 WT and GATA4^ΔIEC^ mice (color, as in A), three days after transfer. **** P<0.0001, *** P<0.001, ANOVA with Tukey multiple comparison test. *N*= 5 mice/group. **(F)** Number (y axis) of transferred T cells (as in E) in the jejunum LP(dots) 9 days after transfer into WT and GATA4^ΔIEC^ mice (color, as in A). ** P<0.01, t-test. *N*= 4 mice/group. **(G)** Representative plot showing frequency of cells with vβ14+ TCR among transferred CD4+ CD45.1+ 7B8+ T cells in in the jejunum of WT (top) or GATA4^ΔIEC^ (bottom) mice. **(H)** Rat SFB loads (y axis), as measured by qPCR relative to host DNA, in the jejunum (dots) of monocolonized WT and GATA4^ΔIEC^ mice (color, as in A). (**I** and **J**) Frequencies (y axis) of IFNγ^+^ cells among CD8αβ T cells (I) and of IL-17^+^ cells among CD4 T cells (J) in jejunum (dots) of GATA4^ΔIEC^ mice monocolonized with rat or mouse SFB (x axis). **** P<0.0001, t-test. *N*= 5 mice/group.

Previous studies showed that SFB triggers the upregulation of various adaptive immune pathways (*2, 12–14*). To determine whether jejunal colonization by SFB was responsible for the significant increase in inflammatory T cell immunity observed in the jejunum of SPF GATA4^ΔIEC^ mice (Fig.1 C-E), we performed a series of microbial transfers. First, transplantation of GATA4^ΔIEC^ and littermate control WT microbiota into GF WT and GATA4^ΔIEC^ hosts, respectively, showed that the host genotype determined the immune outcome irrespective of the input microbial community (Figure S2B, C). Furthermore, transfer of either altered Schaedler flora (ASF), a defined bacterial community that lacks SFB, or jejunal microbiota from a WT donor within our colony with low-to-undetectable levels of SFB, failed to induce inflammatory T cells in GATA4^ΔIEC^ mice. These results suggest that SFB is the causative microbe for dysregulated immunity in GATA4^ΔIEC^ mice (Figure S2B, C). Finally, transfer of fecal homogenates from Jackson C57BL/6J mice with or without SFB supplemented demonstrated that the increases in both IFNγ^+^ CD8αβ T cells and T_H_17 cells observed in the jejunum of GATA4^ΔIEC^ mice (Fig. 1D-E), were dependent on the presence of SFB (Fig. 2C, D).

Since SFB colonization is known to induce an antigen-specific T_H_17-cell response against the 3340 epitope of SFB (*15*), we asked if the increased T_H_17 cell response observed in the jejunum was a consequence of altered T cell priming. To address that question, we transferred congenically marked CD45.1^+^ 7B8^+^ CD4 T cells, specific to the 3340 epitope of SFB, into CD45.2 WT and GATA4^ΔIEC^ hosts. In the absence of GATA4, 7B8^+^ CD4 T cells differentiated into RORγt^+^ Foxp3-CD4 T cells in both the jejunal and ileal draining mesenteric lymph nodes (MLN) (Fig. 2E), indicating a change in regional T cell priming against SFB in the MLN. Furthermore, nine days after transfer, 7B8^+^ T cells expanded (Fig. 2F) and downregulated expression of their TCR in the jejunum of GATA4^ΔIEC^, versus WT, mice (Fig. 2G). Together, these results indicate that, through GATA4, IECs regulate regional colonization of the commensal microbe SFB and SFB-specific T-cell priming.

To establish whether colonization of the jejunum by SFB depended on its ability to adhere to IECs, we monocolonized GF WT and GATA4^ΔIEC^ mice with rat SFB, which is 86% genetically identical to mouse SFB but unable to adhere to mouse IECs and thereby induce a T_H_17 response (*12, 16*). Rat SFB was unable to expand and induce inflammatory T cell responses in the jejunum of GATA4^ΔIEC^ mice (Figure 2H-J). To further test the hypothesis that GATA4 prevents colonization of the jejunum by adherent commensals, we transplanted GATA4^ΔIEC^ mice with ASF containing the mucosal-associated commensal microbe *Mucispirillum ASF 457*. Consistent with the hypothesis, ASF colonization of GF GATA4^ΔIEC^ and WT mice led to an expansion *Mucispirillum ASF 457* in the jejunum of GATA4^ΔIEC^ but not WT mice (Fig. S2D).

### GATA4 regulates luminal IgA levels to control colonization by SFB

We next sought to understand how GATA4 in IECs restricts adherent bacterial colonization in the proximal intestine. Expansion of SFB was reported in the ileum and feces of IgA deficient mice (*17, 18*). Furthermore, a previous report revealed that B-cell deficient mice display lipid metabolic defects in the jejunum, as well as a gene expression signature associated with GATA4^ΔIEC^ mice (*19*). Together, these studies suggest the possibility that an IgA defect in GATA4^ΔIEC^ mice controls SFB colonization. In accordance with this hypothesis, SFB monocolonization of GF B-cell deficient Jh^-/-^ mice and GF IgA^-/-^ mice substantially increased SFB loads in the jejunum compared to those observed in the ileum of littermate control mice indicating that IgA regulates regionalization of SFB colonization (Fig. 3A, B, S3A, B).

**Figure 3.**
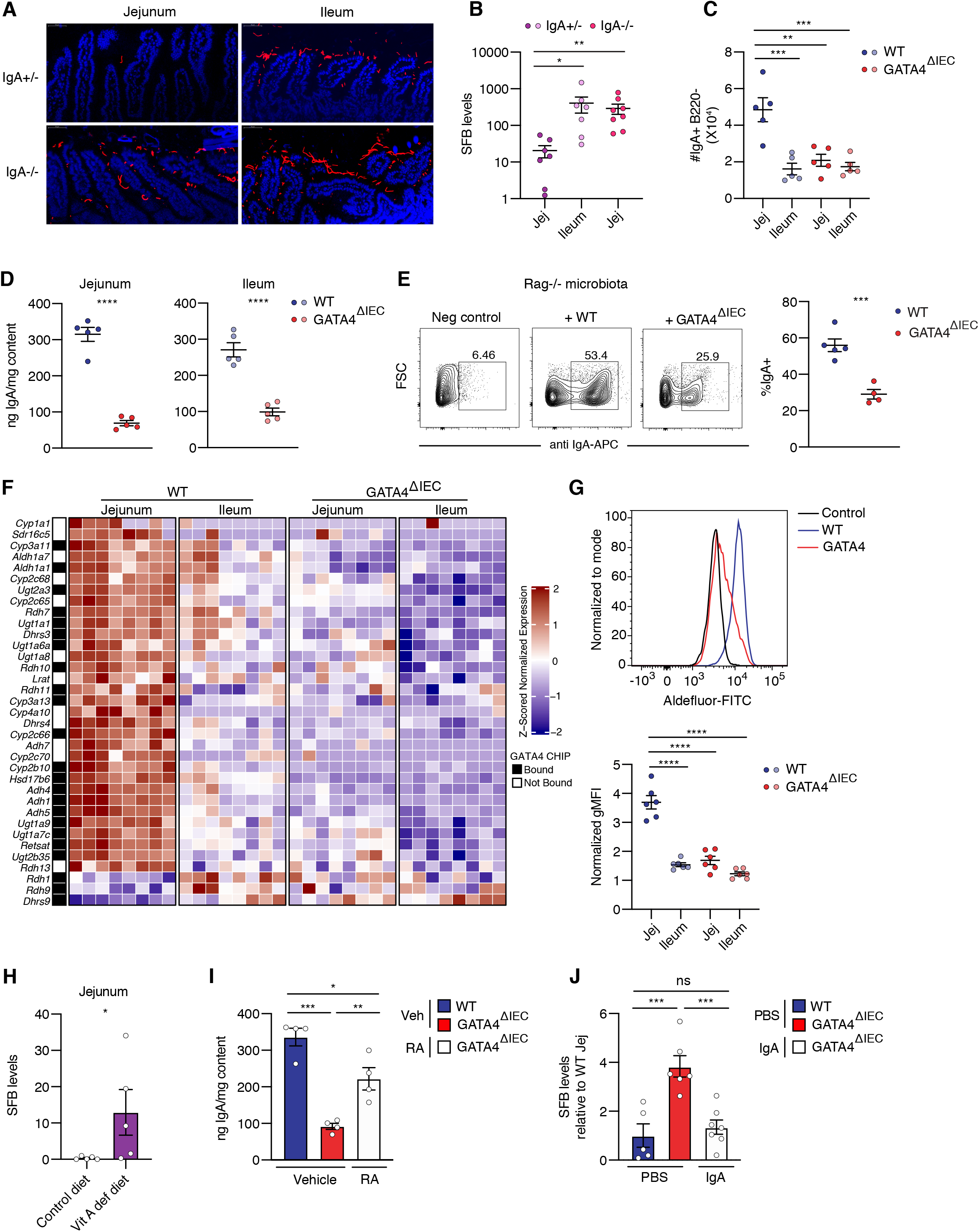
GATA4 regulates regionalization of retinol metabolism and IgA to limit SFB colonization in the proximal intestine. **(A)** FISH staining of SFB from jejunal (left) and ileal (right) tissue of monocolonized IgA deficient (IgA-/-) (bottom) and littermate control (IgA+/^-^) (top) mice. SFB is stained red with Cy5-conjugated SFB-specific 16S probes. The DAPI counterstain is blue. **(B)** SFB load (y axis), as measured by qPCR, relative to host DNA in mucosal scrapings (dots) from the jejunum (violet) and ileum (light pink) of control (IgA+/^-^) mice and the jejunum (fuchsia) of IgA deficient (IgA-/-) mice (x axis) mice. ** P<0.01, * P<0.05, Kruskal-Wallis with Dunn multiple comparison test. *N*= 7-8 mice/group. **(C)** Number of CD45^+^ lin^-^ IgA^+^ B220^-^ plasma cells (y axis), as determined by flow cytometry, in tissue (dots) taken from the jejunum (dark colors) and ileum (light colors) of WT (blues) and GATA4^ΔIEC^ (reds) mice. *** P<0.001, ** P<0.01, ANOVA with Tukey multiple comparison test. *N*= 5 mice/group. **(D)** Concentration of IgA (y axis), as determined by enzyme-linked immunoassay (ELISA), in the contents (dots) of the jejunum (left) or ileum (right) of WT and GATA4^ΔIEC^ mice (color, as in C). **** P<0.0001, t-test. **(E)** Frequency of IgA coated bacteria (gated SYTOBC+) after staining of RAG-/- feces with supernatant from WT and GATA4^ΔIEC^ jejunal contents, followed with labelled anti-IgA APC antibody. *** P<0.001, t-test. *N*= 4-5 mice/group. **(F)** Heatmap of z-scored expression (color) of epithelial, region-specific, GATA4-regulated genes (rows) overlapping the KEGG retinol metabolism pathway, determined from RNA-seq of epithelial cell samples (columns) from the jejunum and ileum of WT (left) and GATA4^ΔIEC^ (right) mice. **(G)** Top, representative histogram of ALDH expression in jejunal epithelial cells of WT (blue) and GATA4^ΔIEC^ (red) mice revealed by ALDEFLUOR staining. WT epithelial cells treated with ALDH inhibitor are shown as negative control for background fluorescence. Bottom, summary plots show the normalized geometric mean fluorescence intensity (gMFI, y axis) of ALDEFLUOR staining in samples (dots) of epithelial cells from the jejunum and ileum of WT and GATA4^ΔIEC^ mice (color, as in C). **** P<0.0001, ANOVA with Tukey multiple comparison test. *N*= 6 mice/group. **(H)** SFB loads (y axis), as measured by qPCR relative to host DNA, in jejunal mucosal scrapings of GF WT mice fed a control or vitamin A deficient diet for 4 weeks and subsequently colonized with SFB. * P<0.05 Mann Whitney test N= 5 mice/group. **(I)** Total IgA (y axis) in the jejunal contents of WT, GATA4^ΔIEC^ vehicle-treated, and GATA4^ΔIEC^ RA-treated mice (x axis, color) after 14 days. *** P<0.001, ** P<0.01, * P<0.05, ANOVA with Tukey multiple comparison test. *N*= 4 mice/group. **(J)** SFB loads (y axis), as measured by qPCR relative to host DNA, in jejunal mucosal scrapings in PBS-treated WT or GATA4^ΔIEC^ mice, and IgA-supplemented GATA4^ΔIEC^ mice (x axis, color). *** P<0.001, ANOVA with Tukey multiple comparison test. *N*= 5-7 mice/group.

Given that B cells and IgA prevented SFB from colonizing the jejunum, we asked whether GATA4 may control SFB colonization by regulating the regional distribution of IgA^+^ plasma cells in the small intestine. We observed that the jejunum contained approximately three times as many IgA^+^ B220-plasma cells as the ileum (Fig. 3C), and that the higher numbers of IgA-producing plasma cells were associated with a greater capacity to produce IgA in tissue explants (Fig. S3C). This regionalization of IgA response was GATA4-dependent, as evidenced by significantly reduced numbers of IgA^+^ B220^-^ plasma cells (Fig. 3C) and IgA production in the jejunum of GATA4^ΔIEC^ versus WT mice (Fig. S3C). The overall result was a substantial decrease in luminal secretory IgA in both the jejunum and the ileum of GATA4^ΔIEC^ mice (Fig. 3D). Reduced luminal IgA was also observed in the jejunum and ileum of GF GATA4^ΔIEC^ mice, indicating that the reduction is microbiota independent (Fig. S3D). Moreover, free IgA in the jejunal luminal content from GATA4^ΔIEC^ mice had less capacity than that of littermate-control WT mice to coat the microbiota present in the feces of immune-deficient (RAG^-/-^) mice (Fig. 3E and S3E). Taken together, these results indicate that GATA4 blocks the regionalization of adherent microbes, such as SFB, by increasing luminal IgA.

We hypothesized that epithelial GATA4 mediated IgA levels by controlling region-specific metabolic processes. To identify potential candidates, we performed gene set enrichment analysis in our epithelial RNA-seq data (Fig. S1B) using the KEGG database, which revealed retinol metabolism as a top GATA4-dependent, region-specific pathway (Fig. S3F). Genes in the retinol metabolic pathway were expressed higher in the jejunum of WT, but not GATA4^ΔIEC^, mice, compared to WT ileum (Fig. 3F), supporting previous reports that the proximal intestine facilitates greater vitamin A uptake and metabolism (*20, 21*). Published ChIP-seq data (*10, 11*) shows direct binding of GATA4 to the promoters of 23/35 (66%) of the differentially expressed epithelial genes in the retinol metabolism pathway (Fig. 3F), suggesting direct transcriptional regulation of retinol metabolism in IECs. Concordantly, in the absence of GATA4, jejunal epithelial cells exhibited impaired aldehyde dehydrogenase (ALDH) activity, indicating a decreased capacity to produce retinoic acid (RA) (Fig. 3G).

RA was previously shown to regulate intestinal B cell responses, including upregulating intestinal homing receptors on B cells and IgA class switching (*22*). In addition, epithelial RARα/β reportedly regulates the number of IgA-producing B cells (*23, 24*). Therefore, we asked whether retinol deficiency was sufficient to enable the SFB expansion in the proximal small intestine that we observed in GATA4^ΔIEC^ mice. We fed GF WT mice a control or vitamin-A deficient diet for 4 weeks post-weaning and subsequently colonized them with SFB. SFB colonized the jejunum of mice fed a vitamin-A deficient diet but failed to colonize the mice fed a control diet (Fig 3H). To determine whether the defect in luminal IgA in the small intestine of GATA4^ΔIEC^ mice could be rescued with exogenous RA, we injected GATA4^ΔIEC^ mice intraperitoneally with RA for two weeks. RA significantly restituted luminal IgA in the jejunum of GATA4^ΔIEC^ mice (Fig. 3I). We also tested whether exogenous luminal IgA was sufficient to prevent colonization of the jejunum by SFB in GATA4^ΔIEC^ mice. Polyclonal luminal sIgA, capable of strongly coating microbes from RAG-/- feces, was isolated from the intestinal contents of WT mice with protein L magnetic beads (Figure S3G). This luminal polyclonal IgA or phosphate-buffered saline (PBS) was gavaged to GF WT or GATA4^ΔIEC^ mice prior to and after colonization with SFB (Fig. S3H). This IgA was indeed sufficient to prevent SFB from colonizing the jejunum and ileum of GATA4^ΔIEC^ mice (Fig. 3J, S3I). These data indicate that, by regulating regional retinol metabolism in IECs, GATA4 regulates luminal IgA, which in turn restricts bacterial adhesion to the proximal small intestine.

### Loss of GATA4 control of colonization by an adherent pathogen leads to severe immunopathology

Since GATA4 controlled intestinal colonization by adherent commensal microbes, we tested whether it also regulated colonization by adherent pathogens. *Citrobacter rodentium* is an adherent pathogen that preferentially colonizes the colon but not the small intestine of WT mice (Fig. 4A). In the absence of GATA4, the niche for *C. rodentium* was altered such that the pathogen colonized the small intestine at levels approaching those in the colon of WT mice (Fig. 4A). Furthermore, the colonization in the small intestine of GATA4^ΔIEC^ mice depended on *C. rodentium’s* ability to adhere to IECs. Mutant ΔEAE *C. rodentium*, which lacks the gene intimin required for adherence to IECs, did not colonize the jejunum and ileum of GF GATA4^ΔIEC^ mice beyond levels observed in GF WT mice infected with WT *C. rodentium* (Fig. 4B). These results suggest that GATA4 controls regional colonization of the small intestine not only by commensal bacteria, but also by adherent pathogenic microbes.

**Figure 4.**
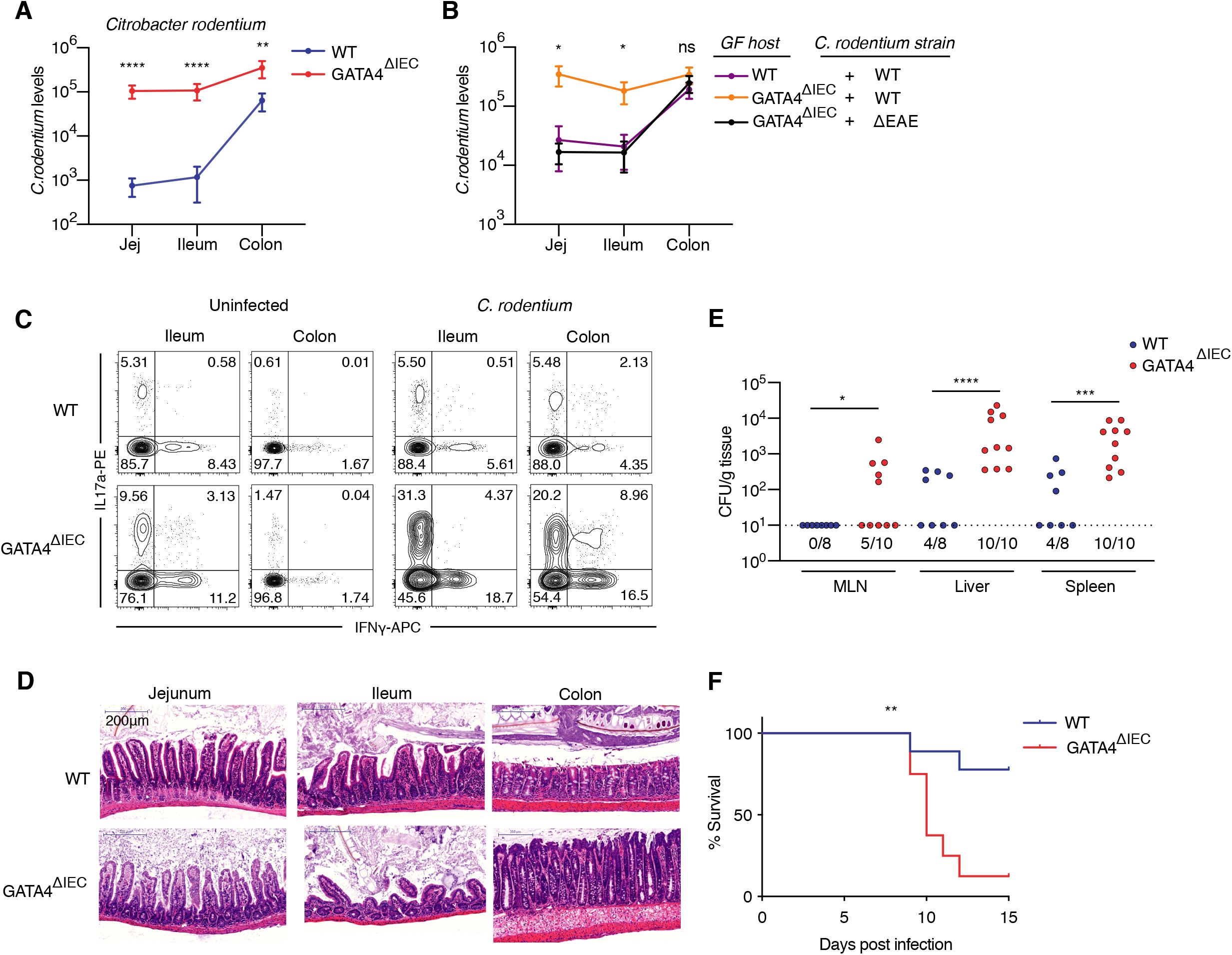
Loss of intestinal regionalization impacts host susceptibility to enteropathogenic infection by *Citrobacter rodentium*. **(A)** *C. rodentium* load (y axis), measured by qPCR relative to host DNA, in distinct intestinal segments (x axis) in WT (blue) and GATA4^ΔIEC^ (red) mice. **** P<0.0001, ** P<0.01, Mann-Whitney test. *N*= 13 mice/group. **(B)** Bacterial loads (y axis) of wild-type *C. rodentium* (purple, orange) and the ΔEAE mutant (black), measured by qPCR relative to host DNA, in distinct intestinal segments (x axis) of GF WT (purple) or GATA4^ΔIEC^ (orange, black) mice. * P<0.05, Kruskal-Wallis with Dunn multiple comparison test, GF WT + WT *C. rodentium* vs. GF GATA4^ΔIEC^ + WT *C. rodentium. N*= 7-9 mice/group. **(C)** Representative FACS plots of IFN-γ (x axis) vs. IL-17a (y axis) in PMA ionomycin stimulated CD4+ T cells that were isolated from the LP of infected or uninfected mice (left) or mice 10 days after infection by *C. rodentium* (right). **(D)** Representative immunohistochemistry (IHC) staining of each intestinal region (panel title) from WT (top) and GATA4^ΔIEC^ (bottom) mice 10 days after *C. rodentium* infection. **(E)** CFUs of *C. rodentium* translocation (y axis) to MLN, liver, and spleen (x axis) 10 days after infection of WT (blue) and GATA4^ΔIEC^ (red) mice. **** P<0.0001, *** P<0.001, * P<0.05, Mann-Whitney test. *N*= 8-10 mice/group. **(F)** Percent survival (y axis) of WT (blue) and GATA4^ΔIEC^ (red) mice 0–15 days (x axis) post *C. rodentium* infection. ** P<0.01 Mantel-Cox test. *N*= 8-9 mice/ group.

We tested whether the expansion of *C. rodentium* colonization to the small intestine of GATA4^ΔIEC^ mice impacted host immune response or survival. In accordance with the change in colonization pattern, *C. rodentium* infection induced a striking expansion of IL-17^+^ IFNγ^-^, IL-17^-^ IFNγ^+^, and IL-17^+^1FNγ^+^ double-positive CD4 T cells in the colon and ileum of GATA4^ΔIEC^ (Fig. 4C). This contrasted with WT littermate control mice, which only showed a moderate T_H_17 and T_H_1 expansion in the colon (Figure 4C). The immune response in GATA4^ΔIEC^ mice was accompanied by the development ten days after infection of severe colitis and villous atrophy in the ileum (Fig. 4D), symptoms not normally associated with the infection in WT. Once the intestinal barrier was compromised in GATA4^ΔIEC^ mice, *C. rodentium* translocated to systemic sites, including the mesenteric lymph nodes, liver, and spleen (Fig. 4E), highlighting the substantial impacts on distal sites that may be triggered by region-specific intestinal defects. By day 12 post-infection, 87.5% of GATA4^ΔIEC^ mice had died (Fig. 4F), punctuating the critical role of GATA4-dependent intestinal regionalization in controlling host disease susceptibility to an enteric pathogen.

### Loss of GATA4 expression in celiac disease is associated with increased IL-17 immunity and changes in bacterial colonization

Celiac disease (CeD) is an auto-immune–like T_H_1-mediated enteropathy of the duodenum caused by dietary gluten in genetically susceptible individuals (*25*). A long-standing conundrum has been the increase of gluten-dependent, but not gluten-specific, duodenal T_H_17 responses in CeD patients (*26–28*). Previous observations noted alterations in IECs (*25, 29*), as well as lipid (*30*) and vitamin deficiencies (*31*) during active CeD (ACeD). We therefore hypothesized that ACeD patients have decreased expression of GATA4 in IECs, and that this decrease may be associated with reported changes in microbiota (*32, 33*) and increased IL-17 immunity. To investigate this possibility, we compared the transcriptional profiles of duodenal biopsies from ACeD patients, CeD patients on a gluten free diet (GFD), and control patients with non-inflammatory intestinal symptoms that required upper endoscopies. Intriguingly, our analysis revealed significantly lower *Gata4* expression in ACeD patients versus controls, which was restored by GFD (Fig. 5A). Immunohistochemistry staining demonstrated an ACeD-associated loss of GATA4 protein production specifically in apical epithelial cells, whereas intestinal crypts cells retained GATA4 production (Fig. 5B). To gain insights into the impact of GATA4 in ACeD, we identified genes that were differentially expressed between “GATA4-hi” individuals, defined by *Gata4* expression above the 70^th^ percentile of the entire cohort, and “GATA4-lo” ACeD patients, defined by *Gata4* expression below the 30^th^ percentile of ACeD patients (Fig. S4A). GATA4-lo specific genes were enriched in mouse genes specific to ileum-like but not jejunum tissue. Furthermore, GATA4-hi specific genes were significantly enriched in mouse genes specific to jejunum but not ileum-like tissue (Fig. 5C, S4B). Thus, during ACeD, intestinal regionalization may be lost as the duodenum decreases GATA4-dependent jejunum-specific gene expression and increases ileum-specific gene expression.

**Figure 5.**
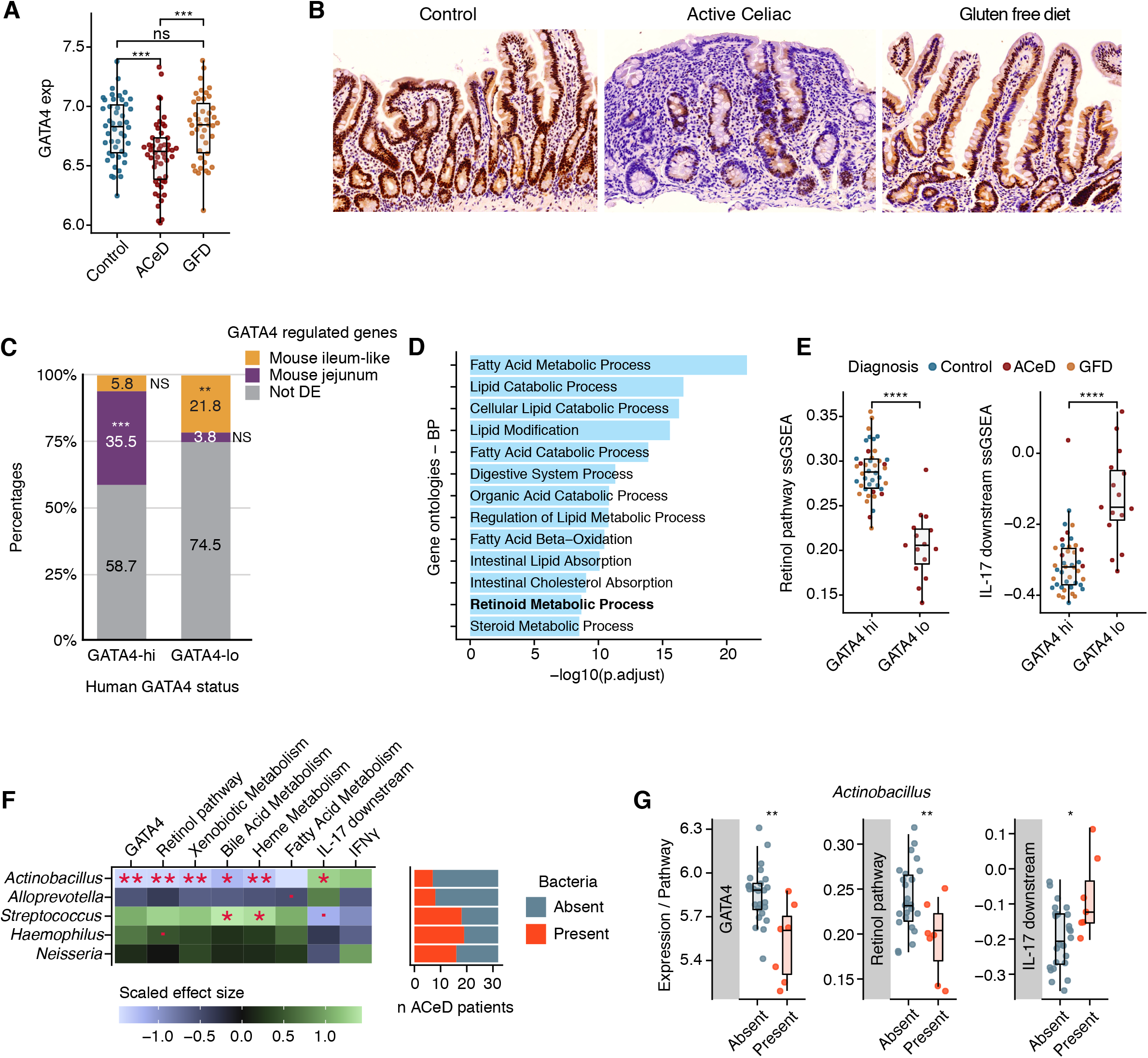
Loss of GATA4 expression is associated with lipid metabolic dysfunction and increased IL-17 signaling in celiac disease. **(A)** Normalized and batch corrected levels of GATA4 expression in duodenal biopsies from healthy controls (blue), active celiac disease patients (ACeD, in red), and gluten free diet celiac patients (GFD, in yellow). *** P<0.001, Wilcoxon test. **(B)** Representative IHC staining for GATA4 in healthy control, active celiac disease patients, and gluten free diet celiac patients. **(C)** Bar plot shows the percentages (y-axis) of human-mouse homologous genes specific to GATA4-hi or GATA4-lo (x-axis) individuals, that are also either jejunum (purple) or ileum-like (yellow) specific and GATA4-regulated, or neither (gray). *** P <10^-51^, ** P <10-6, NS not significant, Fisher’s exact test. **(D)** Bar plots show the pathways (y axis) and significance (–log_10_ of adjusted P, x axis) of the most significantly enriched gene ontologies (biological processes) in the 250 genes that are upregulated in both GATA-hi individuals and WT mouse jejunum. **(E)** Single-sample gene set enrichment analysis (ssGSEA) scores (y axis) for the retinol metabolism (left) and IL-17 downstream signaling (right) pathways in GATA4-hi and GATA4-lo (x axis) individuals (dots) from control (blue), ACeD (red), and GFD (yellow) patient groups. **** P <0.0001 Wilcoxon rank test. **(F)** Left, heatmap displays the scaled effect size (color) of the absence or presence of relevant 5 bacteria (see Fig. S4I) (rows) on GATA4 expression and on the ssGSEA scores of metabolic and immune pathways (columns), while accounting for sex, age and technical covariates. Significance annotated in red, · *P* <0.1, * *P* <0.05, ** *P* <0.01 t-test. Right, bar plot showing the numbers (x axis) of ACeD samples in which the bacteria (y axis) were detected (orange) or not (gray). **(G)** Box plots show GATA4 expression (log_2_ CPM, y axis, left panel) and ssGSEA scores (y axis, other panels) for the retinol metabolism (center) and IL17 downstream signaling (right) pathways in ACeD patients (dots), grouped (x axis) by the absence (blue) or presence (orange) of bacteria from the Actinobacillus genus. * *P* <0.05, Wilcoxon test.

Many of the genes driving these enrichments are direct targets of GATA4 and involved in lipid metabolic processes, such as cholesterol absorption and retinol metabolism (Fig. 5D). Compared to GATA4-hi patients, GATA4-lo ACeD patients demonstrated decreased enrichment in retinol metabolism genes and increased enrichment in IL-17 signaling genes (Fig. 5E). In fact, expression of these pathways was correlated (or anti-correlated, in the case of IL-17 signaling genes) with *GATA4* expression across all ACeD patients (Fig S4C-F). Together, these data reveal that GATA4 expression can be lost in ACeD, and that this loss may play a role in regional defects, such as low retinol metabolism, low plasma cholesterol (*34*), and increased IL-17 immunity.

Studies in mice revealed that the presence of T_H_17 cells in the ileum is dependent on the microbiota (*2*) and that bacteria can drive distinct immune responses depending on the host context (*35*). Therefore, we asked whether a particular microbe was associated with the IL-17 response in ACeD patients. To answer this, we leveraged a quantitative framework to detect absolute abundances of individual bacterial taxa in duodenal biopsies in which host transcriptional analysis is performed concomitantly (*36*). Digital PCR anchoring of 16S rRNA amplicon sequencing revealed no change in overall bacterial load in ACeD patients versus controls (Fig. S4G) and only a trend of stratification between ACeD and control patients in the second and third principal components (Fig. S4I). We did observe an expansion of *Neisseria* in ACeD patients, the only microbial change significantly associated with disease state (Fig. S4H, J), confirming a previous report (*37*). The increased abundance of *Neisseria* was not, however, associated with decreased *Gata4* expression, metabolic defects, or IL-17 signaling in ACeD patients (Fig. 5F).

In contrast, we found that *Actinobacillus* was associated with lower *Gata4* expression, lower retinol metabolism, and higher IL-17 signaling in ACeD, but not control, patients (Fig. 5F, G, S4K). Of note, we did not observe a significant association with the prototypical gluten-specific T_H_1 IFNγ pathway in ACeD patients, suggesting that *Actinobacillus* may play a specific role in IL-17 immunity (Fig. 5F). Intriguingly, *Actinobacillus* was also associated with other metabolic defects in ACeD patients, such as in xenobiotic, bile acid, and heme metabolism, whereas other microbes, such as *Streptococcus*, had inverse patterns and positive associations with bile and heme metabolism. The discovery of mucosal-associated bacteria, such as *Actinobacillus*, associated with the loss of GATA4, metabolic dysfunction, and IL-17 immunity in ACeD patients, highlights the potential importance in the pathogenesis of celiac disease of GATA4 regulation of host microbial interactions.

## Discussion

In this report, we identify mechanisms controlling regionalization of the proximal and distal small intestine and reveal the importance of this segregation in homeostasis and disease. We found that in part by regulating retinol metabolism and local IgA response, the epithelial transcription factor GATA4 plays a critical role in limiting bacterial adhesion in the jejunum, which may allow the host to focus on digestion and absorption of nutrients. On the other hand, the ileum, lacking GATA4 expression and producing less IgA permits adherent microbes, such as SFB, to induce inflammatory IL-17^+^ T cell responses required for controlling pathogenic infections (*2*). The intriguing association between loss of GATA4 expression in IEC and microbial-associated dysregulated IL-17 immune responses in active CeD, poses the intriguing question of whether a decrease in GATA4 in IECs of the proximal intestine may play a role in other complex immune disorders by altering host-microbial interactions and triggering dysregulated immune responses to bacteria.

## Figure legends

**Figure 1 supplemental.**
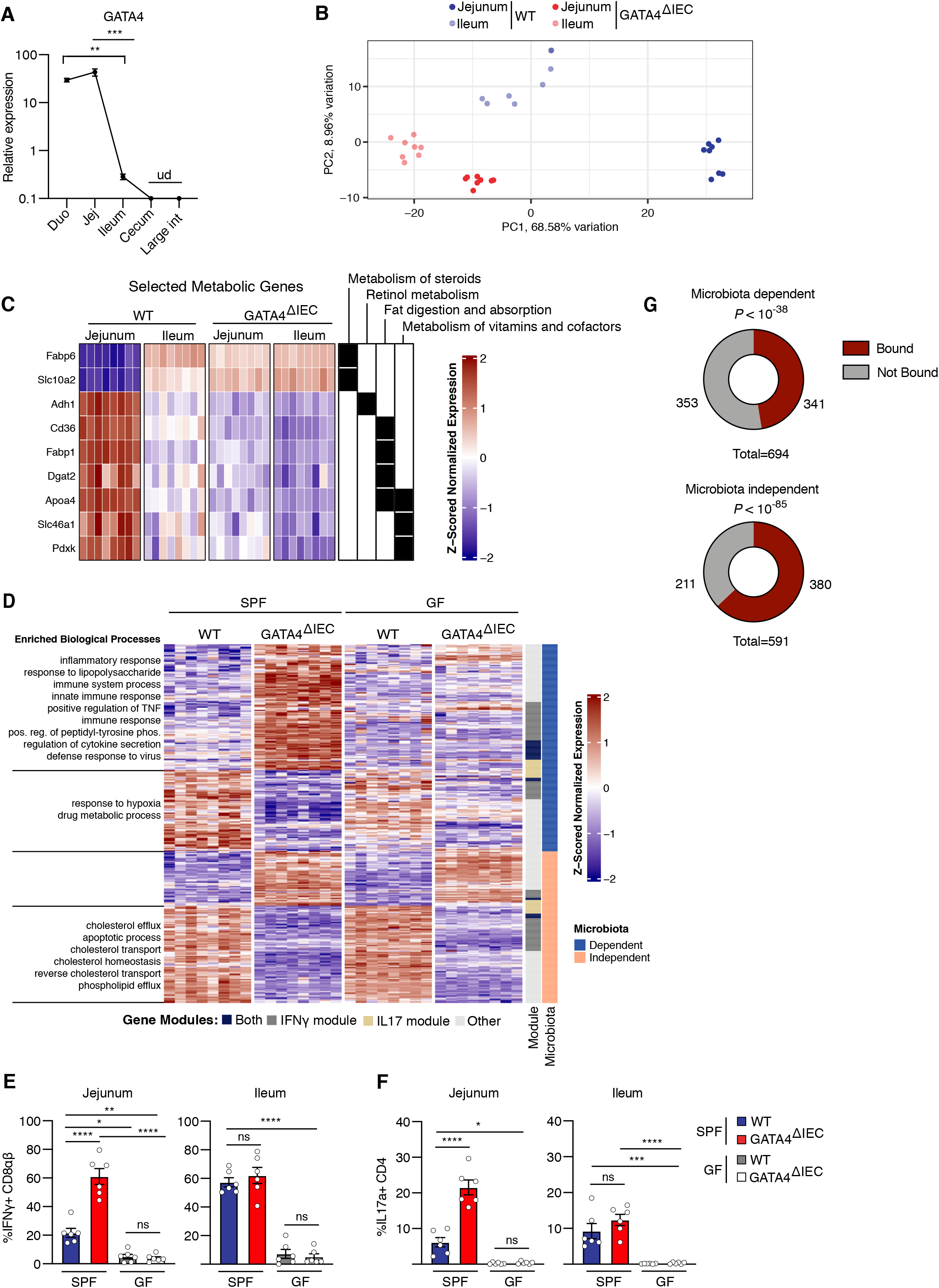
GATA4 controls regionalization of tissue metabolism and immunity. **(A)** Expression of GATA4 (y axis), as measured by qPCR relative to GAPDH, in the tissue of all intestinal regions (x axis). **(B)** Top two principal components (x, y axes) of the expression of region-specific, GATA4-regulated genes in samples (dots) of sorted Epcam^+^ CD45^-^ IECs isolated from the jejunum (dark colors) and ileum (light colors) of WT (blues) and GATA4^ΔIEC^ (reds) mice. **(C)** Heatmap of z-scored expression (color) in IECs of the jejunum and ileum of WT and GATA4^ΔIEC^ mice (columns) of selected DE genes (rows) from the following metabolic pathways: metabolism of steroids (Reactome), retinol metabolism (KEGG), fat digestion and absorption (KEGG), and metabolism of vitamins and cofactors (Reactome). **(D)** Heatmap of the z-scored expression (color) of all region-specific, GATA4-regulated genes (rows) in jejunum tissue (columns) of SPF (left set) and GF (right set) WT and GATA4^ΔIEC^ mice. (**E** and **F**) Frequency (y axis) of IFNγ^+^ cells among CD8αβ T cells (E) or of IL-17a^+^ cells among CD4 T cells (F) in the jejunum (left panel) and ileum (right panel) of SPF and GF WT and GATA4^ΔIEC^ mice (bar color). ****P<0.0001, *** P<0.001, ** P<0.01, * P<0.05, ANOVA with Tukey multiple comparison test *N*= 6 mice/group. **(G)** Frequency of genes bound (red) or not bound (gray) by GATA4, as determined by GATA4 ChIP-seq, among microbiota dependent (top) or microbiota independent (bottom) genes, as in D.

**Figure 2 supplemental.**
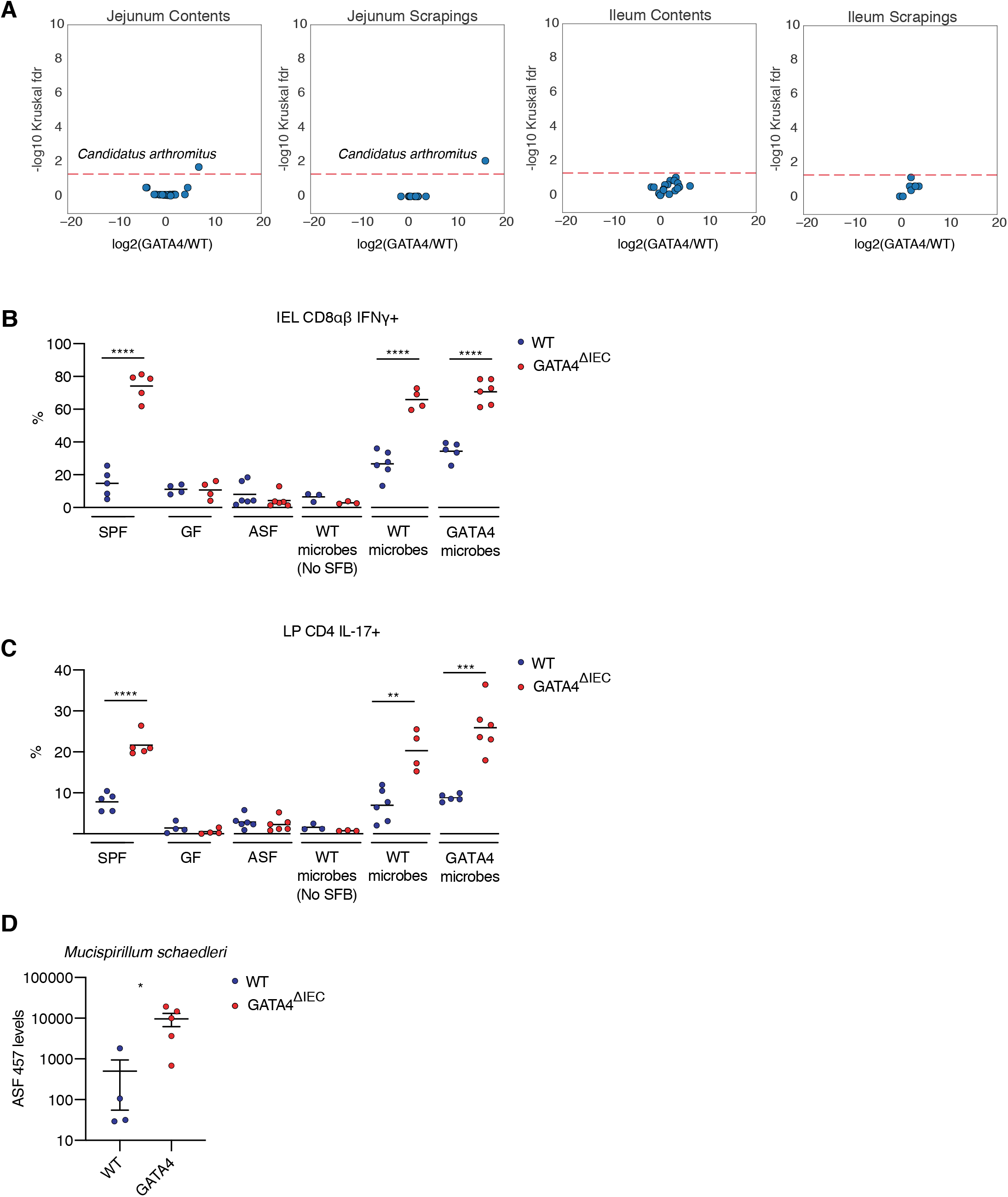
GATA4 controls the colonization of regional mucosal associated bacteria to shape tissue immunity. **(A)** Relative frequencies (x axis, log_2_ fold change) in GATA4^ΔIEC^ versus WT mice of different types of microbes (dots), and their statistical significance (y axis, –log_10_ of the FDR-adjusted *P* value), based on 16S rRNA sequencing of the luminal contents and mucosal scrapings of the jejunum (left set) and ileum (right set) of WT and GATA4^ΔIEC^ mice. SFB are classified as *Candidatus arthromitus*. (**B** and **C**) Frequencies (y axis) of IFNγ^+^ cells among CD8αβ T cells (B) and of IL-17a^+^ cells among CD4 T cells (C) in the jejunum (dots) of WT (blue) and GATA4^ΔIEC^ (red) mice with different microbiota (x axis), *i.e.*, SPF, GF, or ASF, WT microbiota (no SFB), WT microbiota (with SFB), GATA4^ΔIEC^ microbiota, or SFB. **** P<0.0001, *** P<0.001, ** P<0.01, t-test. *N*= 3-6 mice/group. **(D)** *Mucispirillum schaedleri* load (y axis), as measured by qPCR, relative to host DNA in the jejunum after ASF transfer to WT (blue) and GATA4^ΔIEC^ (red) mice. * P<0.05, Mann-Whitney. *N*=4-5 mice/group.

**Figure 3 Supplemental.**
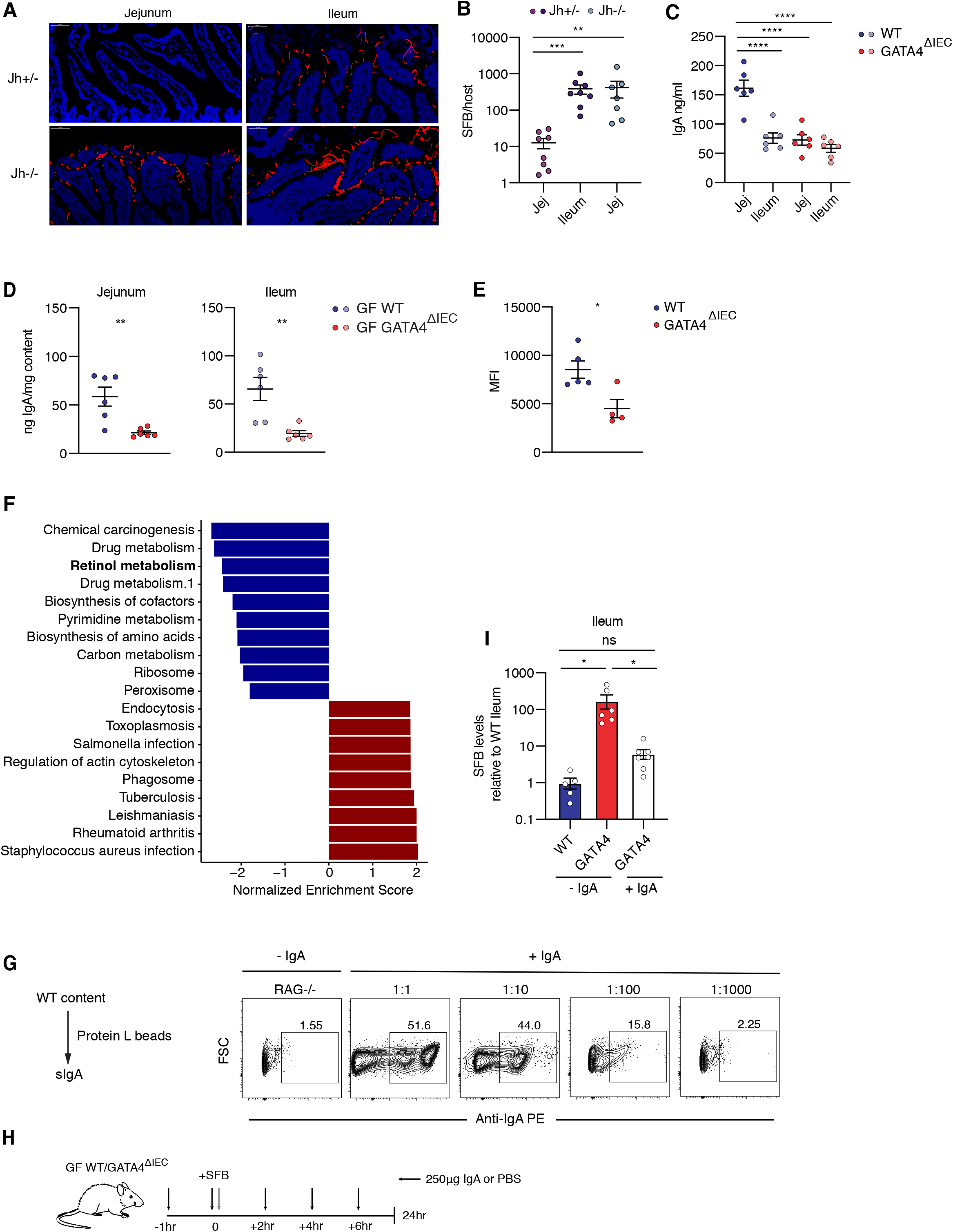
GATA4 regulates regionalization of retinol metabolism and B cell responses. **(A)** FISH staining of SFB in jejunal (left) and ileal (right) tissue of monocolonized B-cell deficient (Jh-/-) and littermate control (Jh+/-) mice. Red indicates SFB stained with Cy5-conjugated SFB-specific 16s probes. Blue is DAPI counterstain. **(B)** SFB load (y axis), as measured by qPCR relative to host DNA, in mucosal scrapings (dots) from the jejunum (light blue) of control (Jh+/-) mice and the jejunum (violet) and ileum (dark purple) of B-cell deficient (Jh-/-) mice. *** P<0.001, ** P<0.01, Kruskal-Wallis with Dunn multiple comparison test. *N*= 7-8 mice/group. **(C)** Concentration of IgA (y axis)), as measured by ELISA, in culture supernatant (dots) of tissue explants from the jejunum (dark colors) and ileum (light colors) of WT (blues) and GATA4^ΔIEC^ (reds) mice after 24 hours in culture. **** P<0.0001, ANOVA with Tukey multiple comparison test. *N*= 6 mice/group. **(D)** Concentration of IgA (y axis), as measured by ELISA, in the contents (dots) of the jejunum (left) or ileum (right) of GF WT (blues) and GATA4^ΔIEC^ (reds) mice. ** p<0.01, t-test. **(E)** Intensity (y axis), of IgA bacterial coating as described in Fig. 3E. MFI, mean fluorescence intensity. * P<0.05, t-test. *N*= 4-5 mice/group. **(F)** Pathways (y axis) in the KEGG database that are significantly enriched (enrichment score, x axis) in epithelial, region-specific, GATA4-regulated genes. **(G)** Schema (left) for isolation of sIgA from luminal contents of WT mice, and FACS plots (right) showing the frequency of IgA-coated bacteria after staining feces of RAG-/- mice with isolated sIgA. **(H)** Experimental schema of IgA gavage and SFB colonization experiment in GF WT and GATA4^ΔIEC^ mice. **(I)** SFB loads (y axis), as measured by qPCR, in ileal mucosal scrapings (dots) in PBS-treated WT or GATA4^ΔIEC^ mice, and in IgA-supplemented GATA4^ΔIEC^ mice (x axis, color). * P<0.05, ANOVA with Tukey multiple comparison test. *N*= 5-7 mice/group.

**Figure 4 supplemental.**
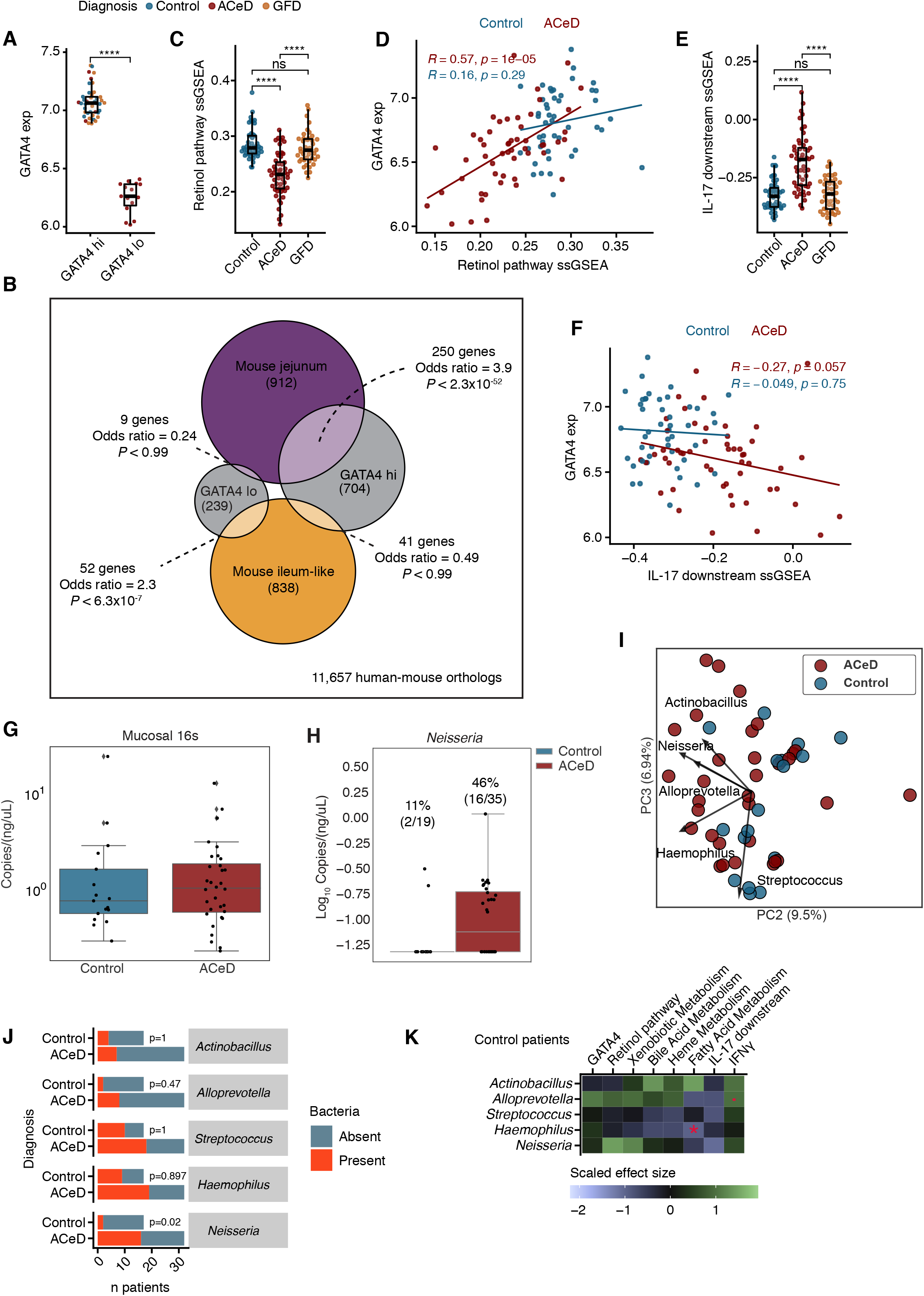
Loss of GATA4 is associated with lipid metabolic defects, mucosal associated bacteria, and increased IL-17 signaling in celiac disease patients. **(A)** *Gata4* expression (y axis) in GATA4-hi and GATA4-lo groups (x axis) in individuals (dots) from control (blue), ACeD (red), and GFD (orange) patients. **(B)** Euler diagram represents the overlaps of 4 sets in a common universe of 11,657 mouse-human transcriptional homologs: GATA4-regulated jejunum-specific (purple) and ileum-like–specific (yellow) genes, and GATA4-hi specific (right, gray) and GATA4-lo ACeD specific genes (left, gray). Enrichment significance as annotated, Fisher’s exact test. **(C)** Retinol pathway ssGSEA scores (y axis) in individuals (dots) from control, ACeD, or GFD patient groups (x axis, color, as in A). **** *P* < 0.0001, NS not significant, Wilcoxon rank test. **(D)** Scatter plot of GATA4 normalized expression (y axis) and ssGSEA scores (x axis) for the retinol metabolism pathway in individual samples (dots) from control (blue) and ACeD (red) patient groups. Annotated with Pearson correlation coefficient and p-value. **(E)** IL-17 downstream signaling pathway ssGSEA scores, analogous to C. **(F)** Scatter plot of GATA4 normalized expression (y axis) and ssGSEA scores (x axis) for the IL-17 downstream signaling pathway, analogous to D. **(G** and **H)** Box plots of absolute abundances (y axis) of mucosal 16S **(G)** and Neisseria **(H)** in biopsies (dots) from control (blue) or ACeD (red) (x axis) patients. **(I)** Principal component analysis (PCA) biplot shows PC2 (x axis) and PC3 (y axis) of each individual (dot) from control (blue) or ACeD (red) patient groups; arrows indicate the loadings of the 5 bacteria most associated with PC2 and PC3. **(J)** Numbers (x axis) of control or ACeD patients (y axis) with detectable (orange) or undetectable (gray) levels of the indicated bacteria. **(K)** Heatmap shows the scaled effect size (color) of each of the bacteria from J (x axis) on GATA4 expression and ssGSEA scores of the indicated pathways in controls, compared to ACeD patients. ≥ *P* <0.1, * *P* <0.05, t-test.

## Materials and Methods

### Mice

7-12 week old mice were used for experiments, co-housed in specific pathogen-free conditions, and kept *Helicobacter hepaticus*, murine norovirus free at the University of Chicago. Some mice were also housed in gnotobiotic isolators and routinely checked for sterility by culture and 16S PCR or kept SFB monocolonized at the University of Chicago Gnotobiotic Research Animal Facility. GATA4fl/fl villin-cre SPF mice were previously generated in the CD1 background and obtained from the Matzinger laboratory (*19*).This line was rederived GF for this study and backcrossed for 10 generations to C57BL/6J background for T cell transfers. C57BL/6J, B6-Tg(Tcra, Tcrb)2Litt/J SFB TCRtg, B6.SJL-Ptprc^a^ Pepc^b^/BoyJ, B.6129S7-Rag1^tmlmom^/J were obtained from the Jackson Laboratory. CD-1 IGS mice were obtained from Charles River Laboratories. B cell deficient mice deficient for IgH J segment locus (Jh) were obtained from Dr. Bendelac at the University of Chicago and generated on a C57BL/6 background using Cas9 with the protospacers GCTACTGGTACTTCGATGTC and GCCATTCTTACCTGAGGAGA. IgA deficient mice where the Sα (IgA switch region) and C1α (first exon) were deleted were obtained from Dr. Bendelac and generated on a C57BL/6 background using Cas9 with the protospacers AAGCGGCCACAACGTGGAGG and TCAAGTGACCCAGTGATAAT. Jh and IgA deficient mice were rederived GF at Taconic Biosciences. Littermate controls of GATA4, Jh, and IgA were used for all experiments in this study. Mice were fed a standard chow diet, vitamin A control diet (Harlan TD.91280), or vitamin A deficient diet (Harlan TD. 86143). Animal husbandry and experimental procedures were performed in accordance with Public Health Service policy and approved by the University of Chicago Institutional Animal Care and Use Committees.

### Patients

A duodenal biopsy was obtained from 166 individuals undergoing upper gastrointestinal endoscopy at the University of Chicago and at Mayo Clinic as previously reported (*38*). There were 64 control patients, 56 untreated patients with active celiac disease, and 46 patients treated with a gluten free diet. All control patients underwent endoscopies for issues unrelated to celiac disease and had normal intestinal histology, no family history of celiac disease, and no significant levels of anti-TG2 antibodies in the serum. Patients with active celiac disease contained positive anti-TG2 antibodies and small intestinal enteropathies with increased IEL infiltration, crypt hyperplasia, and villous atrophy according to the accepted diagnostic guidelines (*39*). The subjects signed an informed consent as provided by the Institutional Review Board of each institution (IRB-12623B for the University of Chicago, and IRB-1491-03 for the Mayo Clinic). DNA and RNA were isolated from each biopsy as described previously (*38*) using the AllPrep DNA/RNA mini kit (Qiagen).

### Isolation of intestinal epithelial cells (IEC), intraepithelial lymphocytes (IEL), and lamina propria (LP) cells

The segments of the intestine were excised as follows to isolate cells for flow cytometry: duodenum was taken 12 cm from the stomach, jejunum 12 cm from the middle, and ileum 12 cm from the cecum. Any leftover segments were discarded. The entire colon was taken after the cecum to the rectum. To isolate IEL and IECs, Peyer’s patches were first removed from the small intestine, and then the segments were opened longitudinally and washed briefly in phosphate-buffered saline (PBS). Epithelial cells, including IELs and LP cells, were isolated as previously described (*40*) using ethylenediaminetetraacetic acid (EDTA) containing calcium-free media and collagenase VIII (Sigma-Aldrich, C2139), respectively. The IEL and LP compartments were then subjected to a 40% percoll density gradient centrifugation step to remove dead cells and debris as previously described (*38*). The IEL and LP cells were then counted on a hemocytometer.

### Cytokine stimulation

Up to 2×10^6^ cells were collected and resuspended in RPMI 1640 media with 10% fetal bovine serum (FBS) and cultured in 48-well plates in the presence of 750 ng/ml of ionomycin, 50 ng/ml of Phorbol 12-myristate 13-acetate (Sigma-Aldrich), and golgi-stop (BD). The cells were incubated for 2 hours at 37C with 5% CO_2_. After stimulation, the reaction was quenched with ice cold FACS buffer, and the cells were subsequently stained with antibodies for flow cytometry.

### Flow cytometry

The cells were first stained with FC block (CD16/32) to block nonspecific binding and then were stained with dead dye to exclude dead cells (Aqua, ThermoFisher or Zombie NIR, Biolegend) for 15 min at 4 °C, followed by staining with cell surface markers for 20 min at 4 °C. For intracellular cytokine staining, the BD cytofix/cytoperm kit was used, and cells were incubated with the antibodies for 40 min at 4 °C. For intracellular transcription factors, the Foxp3 eBioscience kit was used according to the manufacturer’s instructions. The antibodies used are indicated in Table (1). For ALDH staining of IECs, the ALDEFLUOR kit was used (StemCell Technologies), following the manufacturer’s protocol. All cells were gated FSC, SSC, singlets, and live cells. IECs were gated CD45-EpCAM+. 100,000 IECs from the jejunum or ileum were sorted with Aria Fusion (BD Biosciences) into RLT buffer (Qiagen) with β-mercaptoethanol for downstream sequencing analysis. IgA plasma cells were gated as described previously (*41*), i.e. EpCAM^-^, CD45+/dim, lineage negative (Ter119, F4/80, CD3, Ly6G, NK1.1, CD19), IgA+, B220-. CD8αβ IELs were gated TCRβ+, CD4-, CD8α+, CD8β+. CD4 LP T cells were gated TCRβ+, CD4+, CD8α-. The following antibodies and clones were purchased from Biolegend: CD45 Pacific Blue (30-F11), CD4 BV785 (GK1.5), CD4 BV605 (GK1.5), IL10 PE-Cy7 (JES5-16E3), CD45.1 Pacific Blue (A20), Tbet PE (4B10), CD44 PE-Cy7 (IM7), CD62L PE (MEL-14), Epcam PerCP-Cy5.5 (G8.8), CD19 FITC (1D3/CD19), NK1.1 BV605 (PK136), CD11C BV605 (N418), TER119 BV605 (TER-119), F4/80 BV605 (BM8), CD3ε BV605 (145-2C11), Ly6G BV605 (1A8), B220 PE-Cγ7 (RA3-6B2). The following antibodies and clones were purchased from BD: CD8β BUV395 (H35-17.2), CD8α PerCP-Cy5.5 (53-6.7), NK1.1 PE-CF594 (PK136), TCRβ BUV737 (H57-597), TCRβ BV711 (H57-597), CD3ε BUV737 (145-2C11), IFN-g APC (XMG1.2), CD45.2 BUV395 (104), vβ14 TCR FITC (*14–2*), RORyt BV786 (Q31-37). The following antibodies and clones were purchased from Thermo Fisher: TCRgd FITC (eBioGL3), IL17a PE (ebio17B7), FOXP3 eFluor450 (FJK-16s), FOXP3 FITC (FJK-16s), FOXP3 PE-Cy7 (FJK-16s), IgA PE (mA-6E1). The cells were run on the LSRFortessa X-20 Flow Cytometer (BD Biosciences) or the Cytek Aurora and data were analyzed using FlowJo software (Treestar).

**Table 1.**
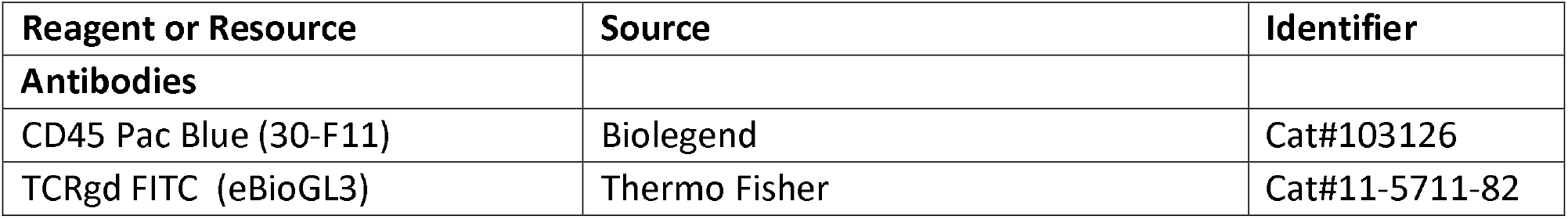
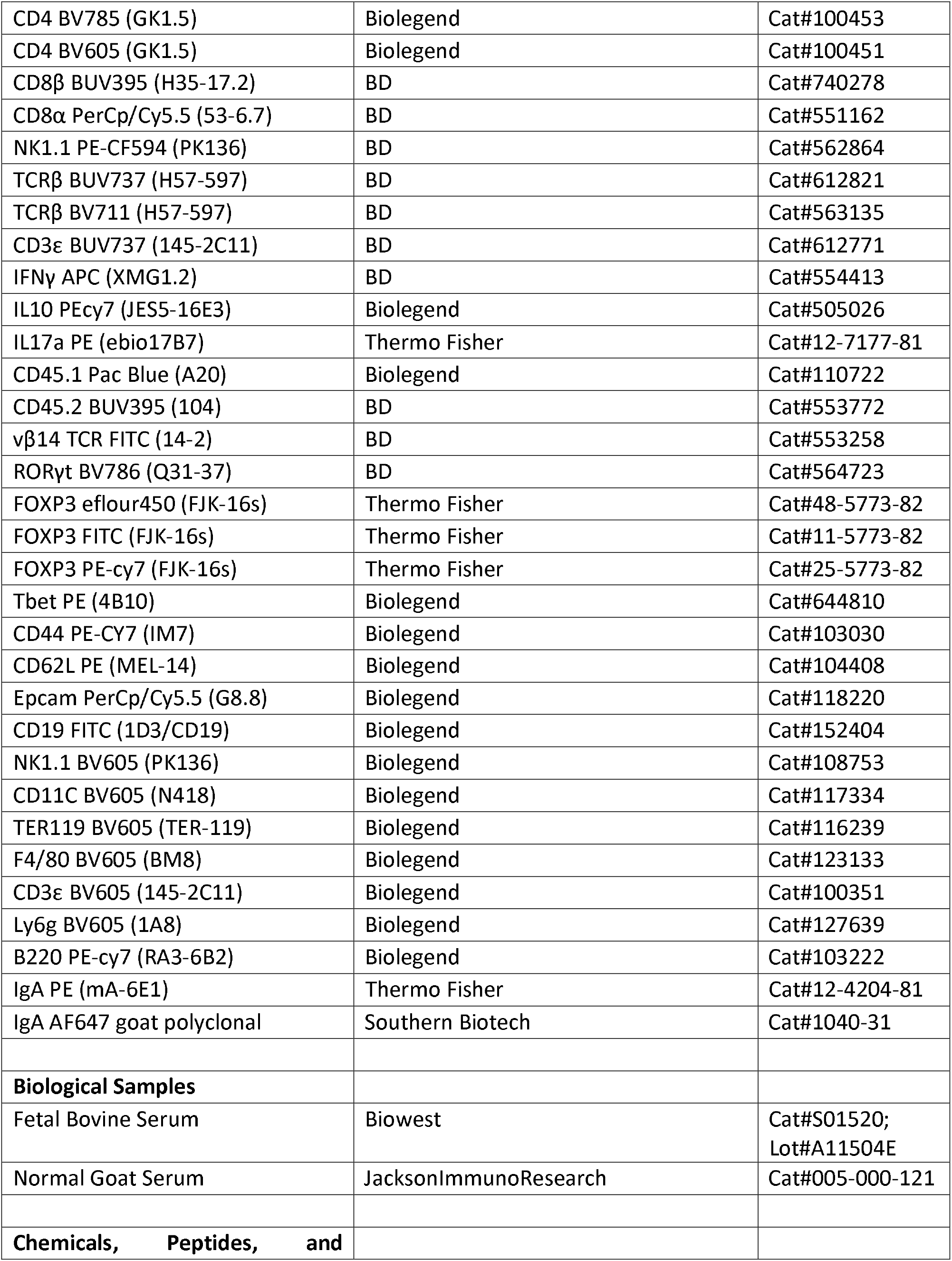
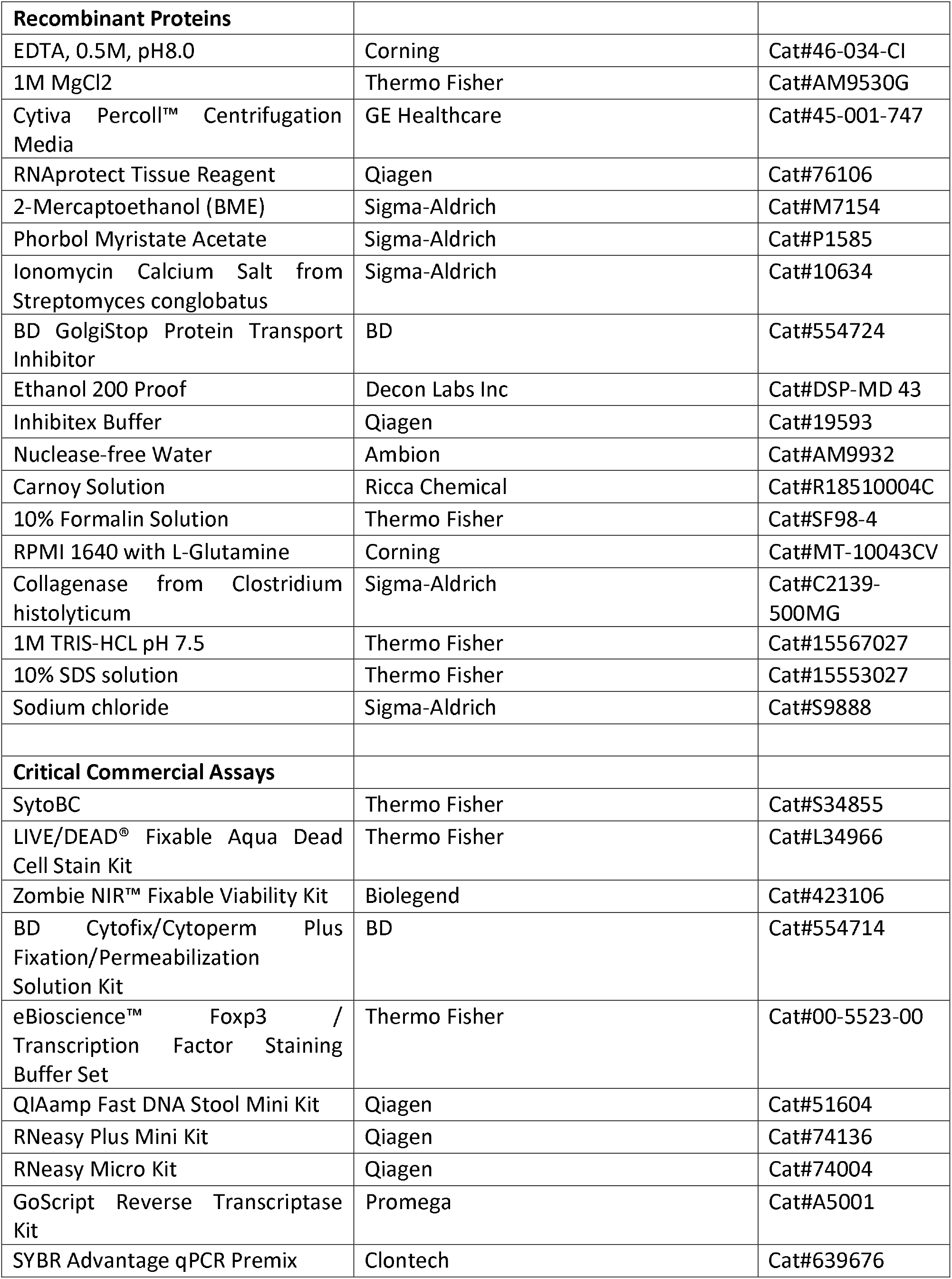
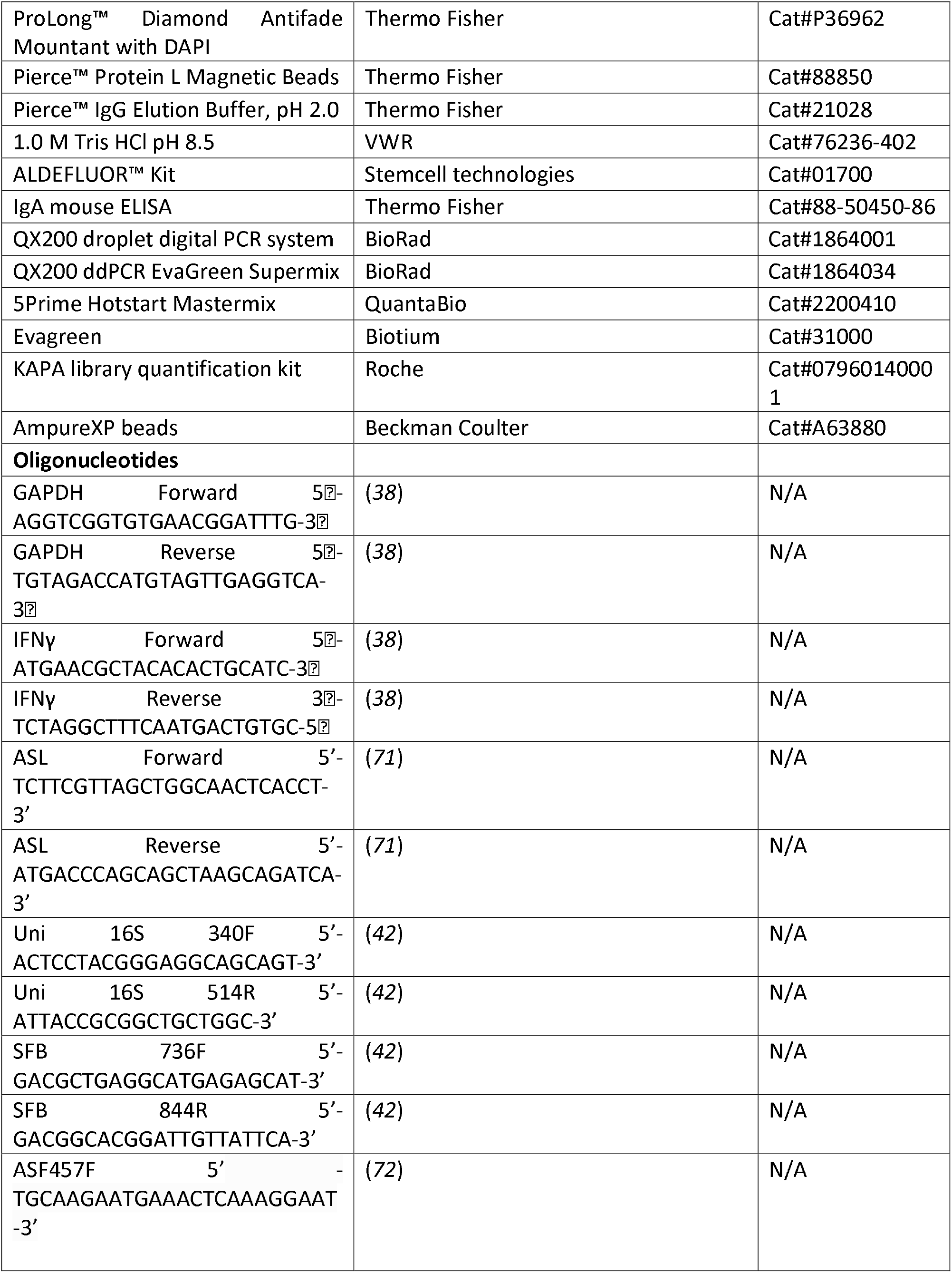
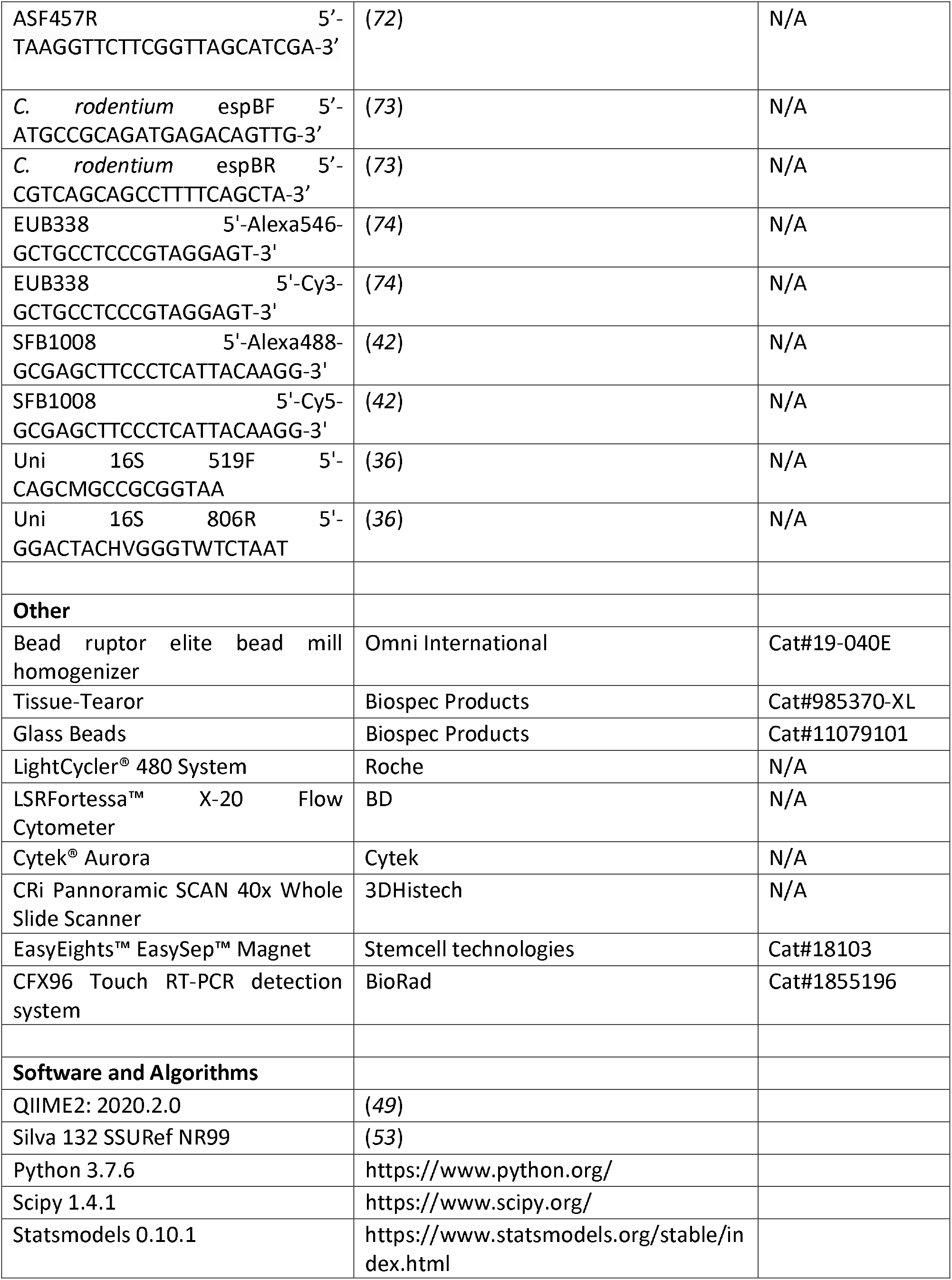
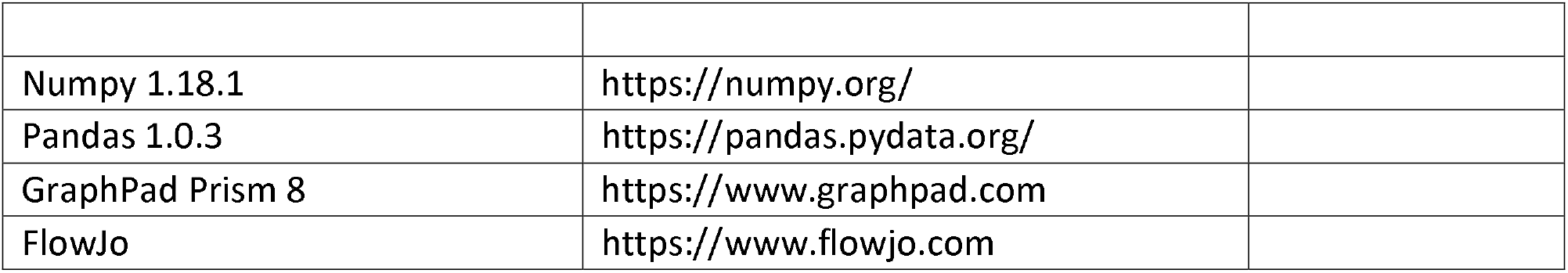

### DNA isolation

For mucosal scrapings for DNA isolation, 5 cm of tissue proximal to the middle of the intestine was taken for the jejunum, and 5 cm from the ileocecal valve was taken for the ileum. The entire colon was used for mucosal scrapings. The tissue was excised, opened longitudinally, scraped with a glass slide, transferred to 2 ml screw cap tube containing 0.1 mm glass beads (Bio-spec), and snap frozen on dry ice. For luminal content, 50-100 mg of content was taken from as close to the middle of the jejunum as possible and from the last 5–7 cm of the ileum. Homogenization was performed after adding 1 ml of inhibitex buffer (Qiagen) using the Bead Ruptor Elite bead mill homogenizer (Omni, 19040E) on speed 6 for 3 min. DNA was then extracted using the QIAmp Fast DNA stool mini kit (Qiagen) following the manufacturer’s protocol with the optional high temp (95 °C) lysis step. DNA concentration was determined using the nanodrop UV spectrophotometer (ThermoFisher).

### RNA isolation

For eventual RNA purification, 1 cm of tissue was excised from the beginning of the duodenum, the middle of the jejunum, the end of the ileum, and the center of the colon and preserved in RNAprotect (Qiagen) overnight at 4 °C and then transferred to −80 °C for long-term storage. The tissue was transferred to 600 μl of RLT buffer containing β-mercaptoethanol (Qiagen) and homogenized for 30 sec with a hand held rotor (Tissue-Tearor, BioSpec). RNA was purified using RNAeasy plus mini kit (Qiagen) following the manufacturer’s protocol with the optional on column DNase digest (Qiagen).

### qPCR

RNA was first reverse-transcribed to cDNA using GoScript Reverse Transcriptase kit (Promega) following the manufacturers protocol. For qPCR, 10 ng of cDNA or 20 ng of DNA from mucosal scrapings and content was used. TB green Advantage qPCR Premix (Takara) was used, and the target gene was quantified and normalized to the housekeeping gene as described previously (*38*) using 1000*2^-(Ct target-Ct housekeeping)^ formula. For host gene expression, the target gene was normalized to GAPDH. For bacterial load, the target gene was normalized to either host DNA as described previously (*42*) with primers specific for host argininosuccinate lyase (ASL) gene or universal 16S primers. The qPCR was performed on the LightCycler 480 System (Roche). The primer pairs and DNA sequences are included in Table (1).

### Histology

The tissue was collected in the same manner as for RNA, placed in cassettes and fixed in 10% formalin for H&E staining or Carnoy solution (ThermoFisher) for fluorescent *in situ* hybridization (FISH) staining overnight at room temperature. Cassettes were transferred to 70% ethanol for formalin or 100% ethanol after Carnoy fixation to wash out the fixative. The tissue was embedded in paraffin, and slides were cut at 5 μm thickness. The H&E staining was performed by the Human Tissue Resource Center at the University of Chicago. For FISH staining, the paraffin was first removed by running the slides through four 3-min incubations in xylene and four 3-min incubations in 100% ethanol. The slides were then moved to a polypropylene slide container and filled with hybridization solution containing the diluted 16S probe (0.9M NaCl, 20mM Tris-HCL pH 7.5, 0.1% SDS with 0.2ng of probe specific for SFB 16S or universal 16S) (*43*). The 16S probes used are included in Table (1). The slides were incubated overnight at 50 °C in the dark. The slides were washed three times with the hybridization buffer, briefly rinsed in H_2_0, and then mounted with Prolong diamond antifade with DAPI (ThermoFisher). The slides were scanned with the CRi Pannoramic SCAN 40x Whole Slide Scanner at the University of Chicago Integrated Light Microscopy core.

### ELISA

To quantify luminal IgA levels, content was collected from the jejunum and ileum and weighed in 2-ml bead beating tubes containing 0.1mm glass beads. After adding 1 ml of 1X cell lysis buffer with protease inhibitors (Cell Signaling Technologies), the content was homogenized on a vortex for 5 min. The debris were pelleted at 13000 rpm for 10 min, and the supernatant was collected for ELISA. For tissue explants, 1 cm of tissue was excised and opened longitudinally, washed in PBS, and placed in complete RPMI at 37 °C for 24 h. The culture supernatant was collected and used for ELISA. The supernatant was diluted in 1X assay diluent A (ThermoFisher), and the dilution in the middle of the standard curve was used to quantify IgA levels. IgA mouse uncoated ELISA kit (ThermoFisher) was used following the manufacturer’s protocol, and absorbance was read at 450 nm. The amounts of IgA were back calculated to the original sample and normalized relative to the weight of the content or to ml of culture supernatant.

### Luminal IgA isolation and *in vivo* treatment

To isolate luminal polyclonal sIgA from the intestine, luminal content was pooled from the small intestine, large intestine, and cecum from SFB+ 8–12 week old WT CD1 mice (Charles river). Content was transferred to falcons containing 1X Tris-Buffered Saline, 0.1% Tween 20 (TBST) buffer with proteinase inhibitor (Roche). Falcons were then vortexed for 5 min on max speed and centrifuged for 10 min at 5000 rpm. The supernatant was collected and spun again two times to further remove bacteria and debris. Pierce Protein L Magnetic Beads (Thermo Scientific) were added and incubated for 1 h at room temperature while shaking. After 1 h, beads selectively bound to IgA through kappa light chain were separated from the supernatant with EasySep magnetic stand (StemCell technologies). Supernatant was discarded and beads washed 3 times. IgA was separated from the beads with Pierce IgG Elution Buffer pH 2.0 (ThermoFisher). Elution buffer was incubated with the beads for 10 min at room temperature on a shaker. Tris-HCl 1M pH 8.5 was added to neutralize the solution. IgA protein concentration was measured using NanoDrop. The isolated IgA preparation was then filtered with 0.22 μm sterile syringe filter unit and protease inhibitors were added (ThermoFisher). The IgA preparation was kept up to one week at 4 °C. When IgA were administered to the mice by gavage, the isolated IgA preparation was further concentrated with Amicon Ultra-4 Centrifugal Filter Units (MilliporeSigma) until 250–350μg/0.1ml final concentration was achieved. To optimize the treatment protocol to restore luminal IgA levels *in vivo* to GATA4^ΔIEC^ mice, we first treated RAG-/- mice with 250 μg of the IgA preparation. After 1 h, we assessed the frequency of IgA+ bacteria in small intestinal contents by flow cytometry as described below, and noted 10–20% of bacteria were IgA+ after gavage. The bacterial coating was transient due to intestinal flow and undetectable after two hours. Therefore, continuous IgA gavages were necessary to sustain luminal IgA and bacterial coating in the small intestine. To administer the IgA preparation and analyze SFB colonization, GF, WT and GATA4^ΔIEC^ mice were gavaged with 100 μl of IgA or PBS. After 1 h, the mice were colonized with SFB and gavaged again with IgA or PBS. Three more gavages were performed at 2-h intervals. The mice were euthanized 24 h after the gavage of SFB, and the regionalization of SFB load was assessed in jejunal and ileal mucosal scrapings by qPCR.

### Bacterial staining with luminal IgA

Luminal content was taken from WT, GATA4^ΔIEC^, or RAG-/- mice and resuspended in 1X PBS with protease inhibitors at a concentration of 0.1mg/μl, vortexed for 5 min, and spun at 8000 rpm for 5 min. Three fecal pellets from RAG-/- mice were homogenized and pelleted. The bacterial pellet was resuspended in 50μl PBS and combined with 50μl of luminal supernatant containing IgA. The IgA was incubated with the bacteria for 1 h at 4 °C. The bacteria were then washed, pelleted, and stained with SYTO BC (ThermoFisher) diluted 1:5000 and anti-IgA APC (Southern Biotech) diluted 1:200 for 30 min. Bacteria were gated on FSC, SSC, SYTOBC+, and IgA+.

### *Citrobacter rodentium* infections

*C. rodentium* strains DBS100, DBS120 *pler-lux*, or DBS100 ΔEAE were grown at 37 °C in Luria broth under agitation (*12, 44*). The cultures were diluted 100X and grew to log phase until the OD^600nm^ reached 0.75. For gavage, 200 μl of bacteria were used, which gave a dose of 2.5×10^9^ CFU/mouse. Mice were separated into cages based on genotype for infections, and male mice were used for survival studies. DBS100 or DBS100 ΔEAE strains were given to GF mice and DBS120 *pler-lux* was given to SPF mice. The DBS120 strain has a genomic kanamycin resistance cassette inserted through Tn5. To determine CFUs of DBS120, 2 fecal pellets/mouse were resuspended in 1 ml of PBS and plated on MacConkey agar containing 50 μg/ml of kanamycin. The CFU/mg feces concentration was determined as: (#CFU counted*Dilution factor/(vol plated in ml))/mg feces. To determine the amount of bacterial translocation, the MLN, liver, and spleen were aseptically dissected, weighed, and homogenized with the Tissue-Tearor rotor (BioSpec) in 500 μl of PBS. Then 200 μl of homogenate was plated on MacConkey agar containing 50μg/ml of kanamycin.

### Microbial transfers

To colonize mice with SFB (*44*) or rat SFB (*12*), 3–4 fresh fecal pellets from SFB monocolonized mice were homogenized in 1 ml of PBS, vortexed for 3 min, and spun at 300 g to remove large debris. Then 200 μl of the homogenate were gavaged to recipient mice. When possible, SFB donor pellets were taken from monocolonized Jh or IgA deficient mice, which harbor 10-fold higher levels of SFB. To colonize mice with SFB-free microbiota, C57BL6 mice from Jackson Labs, which lack SFB in the microbiota, were used as donor mice. Small intestinal and cecal contents were pooled for one donor mouse homogenized in PBS, and gavaged to recipient mice with or without SFB supplemented. For WT and GATA4^ΔIEC^ microbiota transfer to GF WT and GATA4^ΔIEC^ hosts, jejunal content was pooled from two donor SPF GATA4^ΔIEC^ mice or littermate WT mice. Colonization of ASF strains (Taconic) was performed as described previously (*45*) and gavaged to recipient WT and GATA4^ΔIEC^ mice. For all microbial transfers, mice were colonized at 4 weeks of age and analyzed at 8 weeks.

### Vitamin A deficient diet

GF C57BL6 mice were placed on control (Harlan TD.91280) or vitamin A deficient (Harlan TD. 86143) diets from 4 to 8 weeks of age. At 8 weeks, mice were monocolonized with SFB for one week as described above, and the amount of SFB in jejunal mucosal scrapings was quantified by qPCR.

### SFB TCRtg adoptive transfer

Naïve SFB TCRtg Vβ8 CD4 T cells were isolated from LNs and spleen of congenically marked CD45.1 Vβ8+/- female mice using the naïve CD4 T cell isolation kit (Miltenyi), and 2×10^5^ cells/100μl mouse were injected retroorbitally into CD45.2 WT and GATA4^ΔIEC^. Three days after transfer, the mice were euthanized to assess T cell priming and activation in the jejunal and ileal draining MLN as described previously (Esterházy et al., 2019). To assess T cell expansion in the LP of the jejunum nine days after transfer, 50,000 cells were injected/mouse.

### 16S sequencing libraries and analysis

The following 16S methods that were used were adapted from a paper currently in press (*46*). Extracted DNA was amplified, barcoded, and sequenced as described previously (*36, 47, 48*). Briefly, amplification of the variable 4 (V4) region of the 16S rRNA gene was performed in 20 uL duplicate reactions: 8 uL of 2.5X 5Prime Hotstart Mastermix (VWR, Radnor, PA, USA), 1 uL of 20X Evagreen (VWR), 2 uL each of 5 uM forward and reverse primers (519F, barcoded 806R, IDT, CoralVille, IA, USA), variable input volumes of extracted DNA template and nuclease free water. Total DNA input (determined by NanoDrop) was limited to 400 ng to prevent inefficient amplification. Amplification reactions were monitored on a CFX96 RT-PCR machine (Bio-Rad Laboratories, Hercules, CA, USA), and samples were removed in the late exponential phase to minimize chimera formation and non-specific amplification (1,4,5). Amplification cycling conditions were as follows: 94 °C for 3 min, up to 50 cycles of 94 °C for 45 s, 54 °C for 60 s, and 72 °C for 90 s. Successfully amplified duplicates were pooled together and quantified with KAPA library quantification kit (Roche, Basel, Switzerland), and then all samples were combined at equimolar concentrations with up to 96 samples per library. AMPureXP beads (Beckman Coulter, Brea, CA, USA) were used to clean up and concentrate libraries before final library quantification with a High Sensitivity D1000 Tapestation Chip (Agilent, Santa Clara, CA, USA). Illumina MiSeq sequencing was performed with a 2×300bp reagent kit by Fulgent Genetics (Temple City, CA, USA). Raw reads were demultiplexed by Fulgent Genetics. Demultiplexed forward and reverse reads were processed with QIIME 2 2020.2 (*49*). Loading of sequence data was performed with the demux plugin followed by quality filtering and denoising with the dada2 plugin (*50*). Dada2 trimming parameters were set to the base pair where the average quality score dropped below thirty. All samples were rarefied to the lowest read depth present in all samples (48,305 reads) to decrease biases from variable sequencing depth across samples (*51*). The q2-feature-classifier was then used to assign taxonomy to amplicon sequence variants (ASV) with the Silva 132 99% OTUs references (*52, 53*). Resulting read count tables were used for downstream analyses in IPython notebooks.

### Absolute abundance

The total microbial load (bacteria and archaea) of each sample and the absolute abundance of each taxon in individual samples was determined as described previously (*36, 47*). Briefly, the Bio-Rad QX200 droplet dPCR system (Bio-Rad Laboratories) was utilized to measure the 16S concentration in each sample with the following reaction components: 1X QX200 EvaGreen Supermix (Bio-Rad), 500 nM forward primer, and 500 nM reverse primer (Uni 16S 519F, Uni 16S 806R) and thermocycling conditions: 95 °C for 5 min, 40 cycles of 95 °C for 30 s, 52 °C for 30 s, and 68 °C for 30 s, followed by a dye stabilization step of 4 °C for 5 min and 90 °C for 5 min. The final concentration of 16S rRNA gene copies in each sample was corrected for dilutions and normalized to the extracted sample total DNA measurement from NanoDrop. We used total DNA levels as a proxy for tissue mass in biopsy samples. For each sample, the input-DNA-normalized total microbial load from dPCR was multiplied by each amplicon sequence variant’s (ASV) relative abundance to determine the absolute abundance of each ASV.

### Poisson quality filtering

Two separate quality-filtering steps based on Poisson statistics were used to determine the statistical confidence in the measured values. First, a 95% confidence interval was calculated from the repeated measures of water blanks. Samples with a total microbial load below the upper bound of this confidence interval were removed from further analysis.

Second, the limit of detection (LOD) in terms of relative abundance was determined for each sample. Sequencing can be divided into two separate Poisson sampling steps. First, an aliquot of sample is taken from the extracted sample and input into the library amplification reaction. The LOD of the library amplification step (LOD_amp_) was determined by multiplying the total microbial load from dPCR by the input volume into the library amplification reaction and then finding the relative abundance corresponding to an input of three copies. Poisson statistics tells us that the likelihood of sampling one or more copies with an average input of three copies is 95%. The second Poisson sampling step in sequencing arises from the number of reads generated from the amplified library. Here, the relative abundance at which 95% confidence of detection is observed was previously shown to follow a negative exponential curve with respect to the total read depth: LOD_seq_ = 7.115 * read_depth^-0.115^ (*36*) The minimum LOD = min(LOD_amp_, LOD_seq_) of the two described LODs was then determined for each sample. For each sample, the abundance of any ASV with a relative abundance below the LOD was set to zero. After this filtering step, data tables for each taxonomic level were generated.

### Statistical analysis and correlations

Group comparisons were analyzed using the non-parametric Kruskal-Wallis rank sum test with Benjamini–Hochberg multiple hypothesis testing correction using *SciPy.stats Kruskal* function and *statsmodels.stats.multitest multipletests* function with the *fdr_bh* option.

### Purified IECs RNA-seq

To perform RNAseq on IECs, 100,000 EPCAM+ CD45-cells were cell sorted from the jejunum and ileum of WT and GATA4^ΔIEC^ mice. Three independent cell sorting experiments were performed and the libraries and sequencing were done on the same batch with 8 mice per group. The SMART-Seq v4 Ultra Low Input RNA Kit (TaKaRa) was used to generated amplified cDNA, using 7500 pg of RNA input. The cDNA was generated and purified according to the manufacturer’s specifications, and cDNA was amplified 12 cycles. The Nextera XT DNA Library Preparation Kit (Illumina) was used to generate the RNA-seq libraries, with an input of 125 pg cDNA, according to the manufacturer’s specifications. Subsequently, the libraries were multiplexed and sequenced at a depth of 20 million reads per sample (50bp SR) on a HiSeq4000.

### Whole tissue RNA-seq

Whole tissue biopsies stored at -80 °C were thawed on ice and transferred to Starstedt tubes containing 350uL RLT Plus supplemented with 1% 2-mercaptoethanol and equal quantities of 1.0 mm and 0.5 mm zirconium oxide beads (Next Advance). Biopsies were bead beat 3 times for 1 min at a setting of 9 on a Bullet Blender 24, with 1 min of cooling on ice between each beating. Lysates were processed using the AllPrep DNA/RNA/miRNA Universal Kit (Qiagen). To generate sample libraries, 500 ng of purified RNA was used as input in the TruSeq Stranded mRNA Library Prep kit (Illumina) according to manufacturer’s specifications. Libraries were multiplexed and sequenced at a depth of 20 million reads per sample (50 bp SR) on a HiSeq4000.

### Mouse RNA-seq data processing to obtain raw counts

Mouse RNA-seq raw data were processed using a standard workflow based on the GENPIPES framework (*54*). Specifically, the “stringtie” type “rnaseq” pipeline was used. Reads were first trimmed using Trimmomatic software (*55*). Trimmed reads were aligned to the Mus_musculus.GRCm38 mouse reference genome using the STAR aligner (*56*) following a two-pass mapping protocol. Alignments were then sorted and filtered for duplicates using Picard(sort, markduplicates) (“Picard Toolkit” Broad Institute http://broadinstitute.github.io/picard/; Broad Institute). Gene-level read counts for downstream processing were calculated from spliced alignments using HTseq count (*57*).

### RNA-seq data analysis: quality control filtering and normalization

All statistical analyses of the mouse RNA-seq data were performed using R(v4.0.3).The raw count matrices contained the transcript counts, with columns representing samples and rows indexed by genes. Genes expressed (*i.e*., having at least two counts) in fewer than two samples were removed. The resulting matrices will be referred to as the count matrices. The variance stabilizing transformation from DESeq2 R package (*58*) was applied to normalize the count matrices. Batch effects were removed using the *removeBatchEffect(*) function from the limma R package (*59*).

### RNA-seq data analysis: differential expression and gene set enrichment

A negative binomial generalized linear model in the DESeq2 package was used to test whether genes were differentially expressed (DE) between sample groups. The analysis takes the count matrix as an input and estimates size factors and dispersions prior to fitting the model. Wald statistics were used to determine the significance and the log2-fold change (LFC) of the fit for each gene. We used the Benjamini-Hochberg method for reducing the false discovery rate (FDR), with a cutoff of <0.05 for identifying differentially expressed genes for further analysis. *P* values reported below are FDR adjusted. The LFCs and FDR-adjusted *P* values were given as input to the *fgsea(*) function from the fgsea R package (*60*), which implements a preranked gene set enrichment analysis. The rankings of the genes were based on the FDR-adjusted *P* values. The mouse KEGG pathway database (mmuKegg) (*61*) and/or the Gene Ontology, Biological Processes (GO-BP) database (*62, 63*) were the gene sets used in the enrichment analysis. Enriched pathways (*i.e., P*<0.05) were collapsed to independent pathways to avoid repetition, using the fgsea *collapsePathways(*) function (*60*).

#### (i) Identifying region-specific GATA4-regulated genes

We determined this set of DE genes by grouping together “ileum-like” samples, *i.e.*, WT ileum and GATA^ΔIEC^ jejunum samples, and comparing them with WT Jejunum samples. This was done separately for the tissue and EC RNA-seq analyses. A threshold of >0.25 for the absolute value of the LFC was used to filter the very high number of DE genes in the tissue RNA-seq data, whereas no LFC threshold was used for the EC data.

#### (ii) Microbiota dependent and independent genes

*We* determined the influence of microbiota on the region-specific GATA4-regulated genes by first systematically comparing jejunum tissue samples to identify: (1) DE genes between GATA^ΔIEC^ SPF in comparison to WT SPF, (2) DE genes between GATA^ΔIEC^ GF in comparison to WT GF, and (3) genes whose expression indicated a significant interaction between genotype (*i.e.*, KO vs WT) and microbiota (*i.e.*, GF vs SPF).

a. Microbiota dependent genes: Genes that were strongly DE (*i.e., P* <0.01, | LFC | > 0.6) in Group 1 but not DE (*i.e., P* >0.2) in Group 2, or vice versa, were deemed microbiota dependent. Further, genes in group 3 that had a strong, significant interaction term (*i.e., P* <0.01, | LFC | > 0.9) were also included.
b. Microbiota independent genes: Genes that were strongly DE (i.e., P <0.01, |LFC| > 0.6) in both groups were deemed microbiota independent. Further, genes in group 3 that also had a weak or insignificant interaction term (i.e., *P* >0.2 or | LFC| < 0.9) were also added to the list of microbiota independent genes.

Both these sets of genes were intersected with the region-specific, GATA4-regulated genes, resulting in region-specific, GATA4-regulated genes that are either microbiota dependent or independent.

#### (iii) Genes regulated by retinoic acid

We determined this set of DE genes by grouping together RA-treated, GATA^ΔIEC^ jejunum and vehicle-treated WT jejunum samples, and comparing them to vehicle-treated GATA^ΔIEC^ jejunum samples. Due to the small number (4) of samples per group, many genes were dropped due to the FDR threshold. To get a better view of the overall trends, DE genes were instead determined in this step by thresholding on the unadjusted p-value (P <0.05). As a sanity check, we observed that using the same *P* value threshold to compare just vehicle-treated WT jejunum samples with vehicle-treated GATA^ΔIEC^ jejunum samples resulted in a set of DE genes similar to the region-specific GATA4-regulated DE genes identified in the first analysis.

### RNA-seq data analysis: data visualization

#### (i) Principal components plots

The principal components analysis (PCA) was done using the *pca(*) function from the PCAtools R package (Blighe K, Lun A, https://github.com/kevinblighe/PCAtools). The top 500 genes selected by highest row variance in the centered (across rows) normalized count matrix were used to calculate the principal components. Using the full set of genes to calculate the PCA shows the same patterns but with a larger spread amongst sample groups.

#### (ii) Heatmaps

Heatmaps were plotted using the *Heatmap*() function from the ComplexHeatmap R package (*64*). The z-scored (i.e., across rows) normalized expression values were used to plot the heatmaps.

### Annotation of GATA4-targeted genes

A previously published and publicly available table of annotated GATA4 ChIP-Seq peaks (*11*) was used to annotate GATA4 targets among the DE genes reported.

### Generation of IL-17 and IFNγ gene modules

We curated an IL-17 and IFNg modules, which consisted of known gene pathways from established databases and publications. The IL-17 module consisted of genes encompassing the following pathways from mmuKEGG: 04657, 04659 and msigDB: m6335, m19422, m298, m300, m461, m460, m39560, m8578, m8581, m8579, m8927, m8928, and the following papers (*12, 18*).

The IFNg module consisted of genes encompassing the following pathways from msigDB: m22085, m5972, m5970, m4551, m9583, m39363, m161, m6305, m6313, m6696, m6695, m6689, m6688, m6523, m6522, m6513, m6512, m1898, m2913, m8662, m8657, m5913, from the Gene Ontology database: GO:0034341, from the Reactome database: R-913531, and the following papers (*65, 66*).

### Human celiac RNA-seq analysis

Adaptors and low-quality bases were trimmed using Trim Galore (v 0.4.4). Resulting reads were then aligned to the human reference sequence Ensembl GRCh38 release 87 using Kallisto (*67*). Next, the derived pseudo counts were normalized into log2 counts per million reads (CPM) using the voom function from the limma package (v3.46.0) (*59*). To evaluate the transcriptomic changes associated to GATA4 de-regulation in the small intestine and in the context of celiac disease we defined two contrast groups: GATA4 low, to reflect loss of regionalization, and GATA4 hi, as a normal jejunum tissue. To define these two groups, we used the normalized expression of GATA4 were then used to rank all control, ACeD and GFD samples to define two contrast groups of samples. First, by grouping the top 30% GATA4 expressing samples into the GATA4 hi group (ACeD n =6, Control n= 18, GFD n= 18) and, second by considering the bottom 30% of GATA4 expression only from ACeD samples, (n=15) which we defined as the GATA4 low group. Next, to define a universe set for the enrichment analysis we generated a set of 11,657 homologous genes from the human and mouse cohort. We then tested transcriptome wide for significant differences in expression across the GATA4 hi and GATA4 low groups of samples, by using a linear model while accounting for sex, age, batch and technical covariates and correcting for multiple testing. Over-enrichment analysis of gene ontologies biological processes were performed using the enrichGO function from the clusterProfiler (v3.0.4) (*68*). All statistical analyses within the human cohort and the overlaps with the mouse GATA4^ΔIEC^ dataset were performed using R (v4.0.3).

Using on the log2(CPM) expression values, we calculated single sample Gene Set Enrichment scores (ssGSEA) for the Retinol pathway, IL17 downstream genes and the hallmark (*69*), using the R Bioconductor Package GSVA (*70*). These ssGSEA scores where then used to test association between GATA hi vs GATA lo contrasts groups, as well as its association with presence or absence of bacteria. Associations where evaluated using a linear model while accounting for sex, age and technical covariates.

## Acknowledgements

We would like to thank the patients and their family members, as well as the University of Chicago Celiac Disease Center, for supporting our research. We would like to thank Drs. Sonia Kupfer, Carol Semrad, Ritu Verma, Hilary Jericho, and Ian Wilson from the Celiac Disease Center for consenting and recruiting patients. We would like to thank Joaquín Sanz Remón for his help with the general transcriptional analysis of celiac disease and control patient biopsies. We thank Drs. Kenya Honda and Gabriel Nuñez for providing rat SFB and ΔEAE *C. rodentium*, respectively. We thank Ivaylo Ivanov for providing SFB TCR transgenic mice and for helpful discussions. We thank Steven Erickson from Dr. Albert Bendelac’s lab for generating using CRISPR-Cas technologies IgA and Jh deficient mice based on previously published mouse models (Chen et al., 1993; Harriman et al., 1999). We thank Betty Theriault, Kristin Kolar, and the entire Gnotobiotic Research Animal Facility at the University of Chicago for their help with the experiments involving gnotobiotic mice. We thank the Human Tissue Resource Center, the Integrated Light Microscopy Core Facility, Flow Cytometry Facility, and the Genomics Facility at the University of Chicago for their technical support. help. We thank Toufic Mayassi and Cezary Ciszewski for the thoughtful discussions and insightful comments. Finally, we thank Valerie Abadie for critical reading of the manuscript. This work was supported by the National Institutes of Health via T32 AI007090 to Z.M.E., T32 GM007281 to D.G.S., U01AI109695 for A. M., U01 AI125250 to A.B., R01 AI144094 to A.B., R01DK103761 to N.S., R01 DK067180 to B.J., and the Digestive Diseases Research Core Center P30 DK42086 at the University of Chicago to B. J.

## Author Contributions

Z.M.E. and B.J. conceived the study and designed the experiments. S.J.R. designed and oversaw computational analysis of the RNA-seq data, with input from Z.M.E., B.J., J.J.S., and R.A.G. N.S., A.M. and P.M, based on their previously published work and studies performed in gnotobiotic mice, put forward the concept that absence of GATA4 in epithelial cells was driving inflammatory immune responses in the jejunum in a microbiota-dependent manner. Z.M.E., W.L., A.K., D.G.S., J.D.E., and J.T.B. performed the experiments. Z.M.E., W.L., J.J.S, R.A.G., J.T.B., and S.J.R. analyzed and interpreted the data. J.J.S. performed computational analysis of mouse RNA-seq data, and R.A.G. performed computational analysis of human RNA-seq data. J.T.B. performed the 16S sequencing and analysis. D.G.S. and V.D. created the RNA-seq libraries, and S.G. performed the DNA alignments. V.D generated data on GATA4 protein expression and performed the immunohistochemistry analysis in the celiac disease cohort. P.M. supervised an initial microarray transcriptional analysis. I.L.T, S.G contributed to and L.B.B supervised RNA-seq analysis of human celiac disease patients. R.F.I. supervised the 16S analysis. A.B. supervised the IgA experiments. Z.M.E, S.J.R, and B.J. wrote the manuscript. W.L., J.J.S, J.T.B., R.F.I., A.M., N.S., and A.B. edited the manuscript. R.F.I., A.M., A.B., N.S., and B.J. acquired funds to support the work; and B.J. directed the study.

